# A Five-Gene Stromal-EMT Signature Predicts Prognosis, Immunotherapy Resistance, and Therapeutic Vulnerability in Bladder Cancer

**DOI:** 10.64898/2026.08.15.745016

**Authors:** Wen Zhang, Shiliang Ji

## Abstract

**Background:** Bladder cancer has entered an era in which immune checkpoint blockade (ICB) and antibody-drug conjugate (ADC)-based combinations are reshaping clinical management. However, transcriptomic scores that connect prognosis, tumor microenvironment state, and treatment response are incompletely defined.

**Methods:** Open-access TCGA-BLCA RNA-seq, clinical, mutation, copy-number, and RPPA data were downloaded from the Genomic Data Commons (GDC). Tumor-normal differential expressions, survival screening, LASSO-Cox modeling, train-test validation, GEO validation, pathway enrichment, immune signature scoring, mutation/CNV/RPPA support, drug sensitivity prediction, single-cell/spatial localization, and ICB validation were performed using reproducible Python and R scripts. A reduced model was derived using only genes shared by TCGA, GSE13507, and GSE31684. The fixed formula was then applied without refitting to IMvigor210 and GSE176307.

**Results:** A five-gene model composed of *EMP1*, *AHNAK*, *TNFRSF14*, *CLEC2D*, and *GSDMB* retained TCGA internal prognostic value (train C-index 0.693, test C-index 0.605, all-sample C-index 0.667; TCGA test log-rank *p* = 0.015), although GEO survival validation in GSE13507 and GSE31684 was modest. High-risk tumors were enriched for epithelial-mesenchymal transition (EMT), TNF-alpha/NF-kB signaling, inflammatory response, hypoxia, complement, CAF, macrophage, checkpoint, and cytotoxic programs. Single-cell and spatial analyses localized the score to basal tumor, endothelial, fibroblast, and perivascular compartments. In IMvigor210, risk scores were higher in ICB non-responders than responders (Wilcoxon *p* = 0.044; AUC for non-response = 0.580), high-risk tumors had a lower responder rate (17.6% vs. 28.0%), and high risk predicted poorer OS (log-rank *p* = 0.016; multivariate continuous risk HR = 3.15, *p* = 0.044). GSE176307 showed directionally consistent but non-significant response results (AUC = 0.576).

**Conclusions:** The five-gene score is best interpreted not as a standalone universal prognostic classifier, but as a compact stromal-EMT and immune-suppression phenotype associated with inferior ICB response. These findings support a framework linking prognosis, microenvironment biology, immunotherapy resistance, and therapeutic hypotheses in bladder cancer.

## Introduction

Bladder cancer is one of the most common malignancies of the urinary tract, which remains a major cause of cancer morbidity and mortality worldwide [1]. Muscle-invasive and metastatic urothelial carcinoma are biologically heterogeneous diseases, with variable clinical outcomes even among patients receiving similar stage-adapted therapy [2, 3]. Over the last decade, large-scale genomic profiling has established that bladder cancer is not a single transcriptional entity. TCGA and related studies have revealed recurrent genomic alterations, luminal and basal differentiation states, stromal and immune programs, and molecular subtypes that carry prognostic and therapeutic implications [4–9]. These observations have motivated efforts to derive gene-expression signatures that move beyond conventional clinicopathologic factors. The treatment landscape of advanced urothelial carcinoma has changed substantially. Platinum-based chemotherapy remains important, but immune checkpoint inhibitors targeting PD-1 or PD-L1 have become central across several disease settings [10–13]. More recently, ADC-based strategies have created a new therapeutic axis. Enfortumab vedotin, an anti-Nectin-4 ADC, has demonstrated survival benefit after platinum and ICB exposure and, in combination with pembrolizumab, has improved outcomes in untreated locally advanced or metastatic urothelial carcinoma [14–16]. Other ADCs, including Trop-2-and HER2-directed agents, have expanded interest in target-antigen expression, payload sensitivity, immune effects, and rational combinations [17, 18]. These advances make it increasingly important to understand whether tumor transcriptomic states identify patients likely to respond to ICB, ADC-based combinations, or other therapeutic strategies.

Although ICB can produce durable responses, only a subset of patients benefits. Prior work using the IMvigor210 cohort showed that response to PD-L1 blockade is shaped not only by tumor mutation burden and T-cell infiltration, but also by TGF-beta-associated fibroblast and immune-excluded phenotypes [19]. More broadly, TGF-beta-driven stromal programs can restrict antitumor immunity and limit checkpoint blockade activity [20, 21]. These findings are particularly relevant in bladder cancer, where basal, mesenchymal, stromal, and immune-infiltrated subtypes overlap in complex ways. A meaningful transcriptomic score should therefore do more than stratify survival, which should help explain tumor microenvironment organization and treatment response. Many published prognostic signatures in bladder cancer are derived from TCGA and validated in GEO cohorts. However, two recurring problems limit translations. Firstly, high-dimensional signatures often include genes that are unavailable across microarray and RNA-seq platforms, making external validation fragile. Secondly, models optimized for overall survival in one cohort may reflect a mixture of tumor-intrinsic biology, tissue composition, treatment history, and platform effects. In our preliminary modeling, a 14-gene TCGA LASSO-Cox model achieved reasonable internal C-index but incomplete GEO gene availability and only modest external validation. We therefore rebuilt a reduced model using only genes shared by TCGA, GSE13507, and GSE31684, accepting a smaller feature set in exchange for cross-platform compatibility.

Here, we present a reproducible multi-omics analysis of TCGA-BLCA and external bladder cancer cohorts. We built a five-gene GEO-compatible risk score, evaluated it in TCGA train-test and external GEO survival settings, interrogated its pathway and immune context, examined mutation/CNV/RPPA support, localized the signature in single-cell and spatial datasets, and validated its correlation with immunotherapy response in IMvigor210 and GSE176307. We further integrated drug sensitivity prediction and discussed how the score may relate to current therapeutic hotspots, including ICB and ADC-based treatment. Our central conclusion is deliberately cautious since the five-gene score should not be framed as a definitive universal prognostic tool. Instead, it provides a compact transcriptomic readout of a stromal-EMT and immune-suppressive state that is biologically coherent and clinically relevant to ICB resistance.

## Materials and Methods

### Study design

This study used public, de-identified datasets and did not require institutional review board (IRB) approval. The analysis was organized into eight linked modules: (1) TCGA-BLCA data acquisition and preprocessing; (2) tumor versus normal differential expression; (3) prognosis-associated gene screening and LASSO-Cox model construction; (4) GEO external survival validation; (5) pathway and immune signature analysis; (6) mutation, CNV, and RPPA support; (7) single-cell and spatial localization; and (8) ICB response validation in IMvigor210 and GSE176307.

### TCGA-BLCA acquisition and preprocessing

Open-access TCGA-BLCA data were downloaded from the Genomic Data Commons (GDC) using project-level API queries [22]. The dataset included STAR gene counts for 431 RNA-seq files, comprising 412 tumor and 19 normal samples, together with clinical survival information for 412 tumor cases, masked somatic mutation MAF files, masked CNV segments, and RPPA protein expression matrices. Gene-level counts were merged into a matrix, transformed into TPM using gene length annotations where available, and summarized at gene-symbol level. For downstream expression modeling, tumor samples were represented as log_2_(TPM + 1). Clinical overall survival (OS) time and event status were derived from vital status, days to death, and days to last follow-up.

### Differential expression analysis

An exploratory tumor versus normal screen was performed using Wilcoxon rank-sum tests on log_2_(TPM + 1) gene-symbol expression. Genes were retained for downstream interpretation when they met false discovery rate (FDR) < 0.05 and absolute log_2_(fold change)>= 1. This screen was treated as exploratory and used primarily to support biological interpretation and figure generation rather than to define the final model alone.

### Prognostic screening and model construction

TCGA tumor samples were split into training and testing sets using stratified sampling by OS event status. A custom Cox-model screening and L_1_-penalized Cox procedure was implemented in Python, inspired by the proportional hazards framework [23] and penalized regression principles [24]. Candidate genes were first screened in the training set. The original 14-gene model was evaluated but showed incomplete cross-platform gene availability in GEO. Therefore, a reduced GEO-compatible model was rebuilt using only genes present in TCGA, GSE13507, and GSE31684. Penalization strength was selected by cross-validation in the training set, targeting a compact 4-8 gene model. The final risk score was fixed as a weighted sum of cohort-wise gene z-scores: Risk score = sum(coefficient_g_ × z(expression_g_)). The final coefficients were *EMP1* = 0.0566, *AHNAK* = 0.0119, *TNFRSF14* = -0.0187, *CLEC2D* = -0.0313, and *GSDMB* = - 0.1004. The formula was not refit in validation datasets.

### External GEO survival validation

GSE13507 and GSE31684 were downloaded from GEO and processed into gene-symbol expression matrices. Probe-to-gene mapping was performed using platform annotation files, and duplicate gene symbols were summarized by average expression. Clinical OS time and event variables were harmonized from available metadata. The fixed five-gene formula was applied using cohort-wise gene z-scores. Survival performance was assessed by Harrell’s C-index [25], median-split Kaplan-Meier analysis, log-rank testing, and Cox regression.

### Pathway, immune, multi-omics, and drug analyses

High-risk and low-risk TCGA samples were compared using median risk score as the threshold. High-versus-low expression differences were ranked and evaluated against MSigDB Hallmark and KEGG gene sets [26, 27]. Enrichment analysis used downloaded gene-set libraries and over-representation tests [28]. Immune and stromal signatures were scored using marker-set averages after gene-wise standardization, conceptually like single-sample gene-set scoring [29]. Somatic mutation MAF files were merged at patient level. Nonsilent mutation burden was calculated per patient, masked CNV segment files were used to estimate burden, and RPPA matrices were compared between high-risk and low-risk samples. In addition to the risk-group burden comparison, raw CNV segment counts were tabulated genome-wide by chromosome across the full TCGA-BLCA cohort to check for chromosome-specific instability hotspots. GDSC2 training data were analyzed using a simplified dual-kernel ridge regression strategy in base R, guided by prior pharmacogenomic response-prediction studies [30–32]. Lower predicted response values were interpreted as greater predicted sensitivity. ADC-specific response was not modeled, because ADC sensitivity depends on target-antigen abundance, internalization, payload biology, and microenvironment context.

### Single-cell, spatial, and immunotherapy validation

Public single-cell and spatial datasets were downloaded and tested for compatibility with the fixed model. GSE145137 primary bladder cancer single-cell RNA-seq data were analyzed using author-provided cell-type annotations [33]. Additional datasets included GSE222315, GSE293189, GSE171351, and GSE269877. GSE222315 and GSE293189 were processed as single-cell datasets using marker-based coarse cell-type inference when author labels were unavailable. GSE171351 was treated as Visium spatial transcriptomics. For each compatible dataset, gene-wise z-scores were multiplied by the fixed five-gene coefficients. For the two candidate single-cell cohorts (GSE222315, GSE293189), gene-level dotplots of the five model genes by inferred cell type and sample-level risk-score distributions stratified by tissue condition were generated to evaluate which model genes drove the cell-compartment pattern and whether the pattern held at the individual-sample level.

IMvigor210 was gained from the official Genentech ‘IMvigor210CoreBiologies’ package. Direct S4 slot extraction was used to export clinical data, selected gene counts, and feature annotations. GSE176307 was downloaded from GEO, which included metastatic urothelial carcinoma patients treated with ICB, with IO response, OS, PFS, *FGFR3* alteration status, TMB, ECOG, sex, and treatment annotations [34]. Response was coded as responder for CR/PR and non-responder for SD/PD. Risk prediction of non-response was evaluated using rank-based AUC, Wilcoxon tests, response-rate comparisons, logistic regression, Kaplan-Meier survival analysis, log-rank testing, and Cox models. To integrate cohort, risk group, immune phenotype or *FGFR3* status, CD8/TGF-beta signature balance, and response into a single view, circos-style association plots and Sankey flow diagrams were generated for IMvigor210 and GSE176307.

### Statistical analysis and reporting principles

All analyses were designed to prioritize reproducibility and conservative interpretation. Continuous variables were summarized using medians or means, and categorical variables were summarized using counts and percentages. For survival endpoints, Kaplan-Meier curves were generated after cohort-specific median risk stratification, and group differences were evaluated using log-rank tests. Cox proportional hazards models were fitted for continuous risk score and for median risk groups where appropriate. Multivariate models were restricted to covariates available in each cohort and were not overloaded relative to event counts. For ICB response, CR and PR were grouped as response, whereas SD and PD were grouped as non-response. Rank-based AUC was used to evaluate risk-score discrimination for non-response, with non-response treated as the positive outcome because high model risk was expected to reflect immune resistance. Wilcoxon tests were used for two-group expression and score comparisons, and Fisher or count-based summaries were used for descriptive categorical associations. Multiple-testing correction was applied to high-dimensional expression screens and enrichment analyses.

TCGA training data were used for feature screening and LASSO-Cox fitting. TCGA testing data, all-TCGA summaries, GEO cohorts, ICB cohorts, and single-cell/spatial datasets were then treated as evaluation or interpretation layers. The five-gene formula was fixed before external application. No coefficient was refit in GEO, IMvigor210, GSE176307, or single-cell/spatial datasets. This design reduces overfitting and makes the external results interpretable. Because this is a retrospective public-data study, all claims were graded by evidence strength. TCGA internal survival, GEO survival, pathway and immune support, single-cell localization, ICB response validation, and drug-response prediction were not treated as equally strong evidence.

## Results

### Study overview and TCGA-BLCA data landscape

The analysis workflow integrated TCGA discovery, GEO validation, multi-omics interpretation, drug prediction, single-cell/spatial localization, and ICB response validation. TCGA-BLCA processing yielded RNA-seq matrices from 431 samples, including 412 tumors and 19 normal bladder tissues. Clinical OS information was available for 412 tumor cases, with 183 OS events used in model evaluation. Mutation, CNV, and RPPA data were available for complementary multi-omics support. External cohorts included GSE13507 and GSE31684 for survival validation, IMvigor210 and GSE176307 for ICB response validation, and multiple single-cell/spatial datasets for cell-compartment localization.

Supplementary Figure S1 documents the TCGA-BLCA data availability matrix, clarifying which samples contributed RNA-seq, clinical, mutation, CNV, and RPPA information to each module. Supplementary Figure S2 adds the AJCC stage distribution for the analyzed TCGA tumors, while Supplementary Figures S3 and S4 provide RNA-seq PCA and tumor/normal sample-count checks. These cohort-context and quality-control panels define the analytic denominator and support the reliability of downstream comparisons.

Exploratory tumor versus normal differential expression identified 2,603 genes meeting FDR < 0.05 and absolute log_2_ fold change >= 1, including 1,056 upregulated and 1,547 downregulated genes in tumor. These genes provided the basis for a broad biological overview but were not used alone to define the prognostic model. The design therefore avoided making a model solely from tumor-normal contrast and instead required survival association, cross-platform gene availability, and downstream biological coherence.

### Reduced GEO-compatible five-gene score retained internal TCGA prognostic performance

The first TCGA LASSO-Cox model contained 14 genes and showed acceptable internal performance, with C-index values of 0.698 in training, 0.614 in testing, and 0.670 across all TCGA samples. However, several model genes were absent from microarray validation platforms, and GEO validation was only modest. To improve portability, we restricted candidate genes to those shared across TCGA, GSE13507, and GSE31684. A reduced model was rebuilt in TCGA training samples and fixed before validation.

The final GEO-compatible model included *EMP1*, *AHNAK*, *TNFRSF14*, *CLEC2D*, and *GSDMB*. *EMP1* and *AHNAK* carried positive coefficients, whereas *TNFRSF14*, *CLEC2D*, and *GSDMB* carried negative coefficients. The fixed formula achieved TCGA train C-index 0.693, test C-index 0.605, and all-sample C-index 0.667. In the TCGA testing set, median-split risk groups showed significant OS separation (log-rank *p* = 0.015), and the test-set multivariate Cox risk-score p value was approximately 0.048. These results indicated that the compact model preserved internal prognostic signal while improving cross-platform feasibility.

Supplementary Figures S5-S10 document the feature-selection path from the exploratory model to the final GEO-compatible score. Supplementary Figures S5 and S6 show the survival separation and risk distribution of the original 14-gene TCGA model, Supplementary Figures S7 and S8 summarize the individual-gene survival screening and Cox volcano landscape, Supplementary Figure S9 reports the original 14-gene coefficient structure, and Supplementary Figure S10 provides the final fixed five-gene coefficients. These panels make the model-reduction process transparent and show why portability across validation platforms was prioritized.

### External survival validation was modest rather than definitive

All five reduced-model genes were available in both GSE13507 and GSE31684. In GSE13507, 165 primary bladder tumor samples with 69 OS events were analyzed. The five-gene score achieved C-index 0.549, but median-split KM analysis was not significant (log-rank *p* = 0.487). In GSE31684, 93 samples with 65 OS events were analyzed. The score achieved C-index 0.547, with non-significant median-split KM *p* = 0.575 (**Figure 1**). These results shaped the interpretation of the study. The five-gene model is compact, platform-compatible, and internally valid in TCGA, but its external survival discrimination in older GEO cohorts is modest. We therefore avoided framing the score as a strong universal prognostic signature and instead asked whether it captured a reproducible biological phenotype with treatment-response relevance.

**Figure 1.**
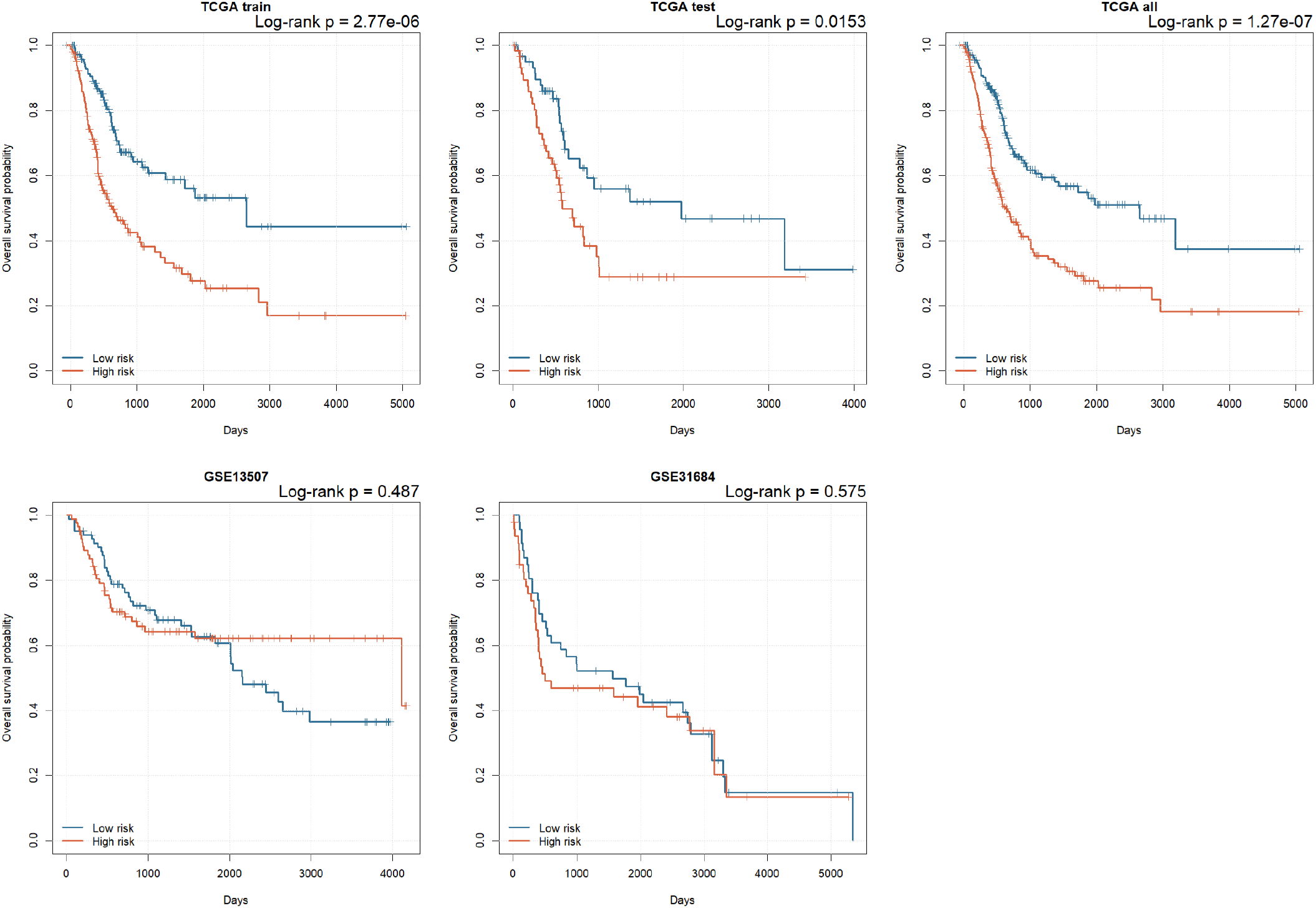
Prognostic stratification of the reduced score across TCGA and GEO validation cohorts.

Supplementary Figures S11 and S12 provide baseline clinical survival context for sex and AJCC stage, helping separate conventional clinicopathologic effects from molecular-score interpretation. Supplementary Figures S13, S14, S15, and S16 further support internal TCGA model evaluation by showing time-dependent ROC curves, KM curves, risk-score distribution, and full-cohort ROC performance for the fixed five-gene score. These supporting panels justify treating TCGA internal prognostic performance as positive but still bounded by external validation.

Supplementary Figures S17-S20 provide detailed GEO validation evidence for GSE13507 and GSE31684. Supplementary Figure S17 shows external KM curves, Supplementary Figures S18 and S20 show risk-score distributions after applying the fixed formula without refitting, and Supplementary Figure S19 shows time-dependent ROC performance. These panels are important because they support cross-platform applicability while also demonstrating the modest external survival discrimination that shaped the cautious interpretation of the model.

### High-risk tumors were enriched for stromal, EMT, inflammatory, and immune-suppressive programs

High-risk TCGA tumors showed marked enrichment of expression programs related to EMT, TNF-alpha/NF-kB signaling, inflammatory response, IFN-gamma response, hypoxia, IL2/STAT5 signaling, complement, and KRAS signaling (**Figure 2**). Immune and stromal marker scoring demonstrated that high-risk tumors had higher CAF, macrophage, M2-like macrophage, checkpoint, cytotoxic/CD8, and inflammatory signatures. These patterns suggested that the risk score reflects an immune-rich but stromal/EMT-dominant tumor microenvironment rather than a simple absence of immune cells.

**Figure 2.**
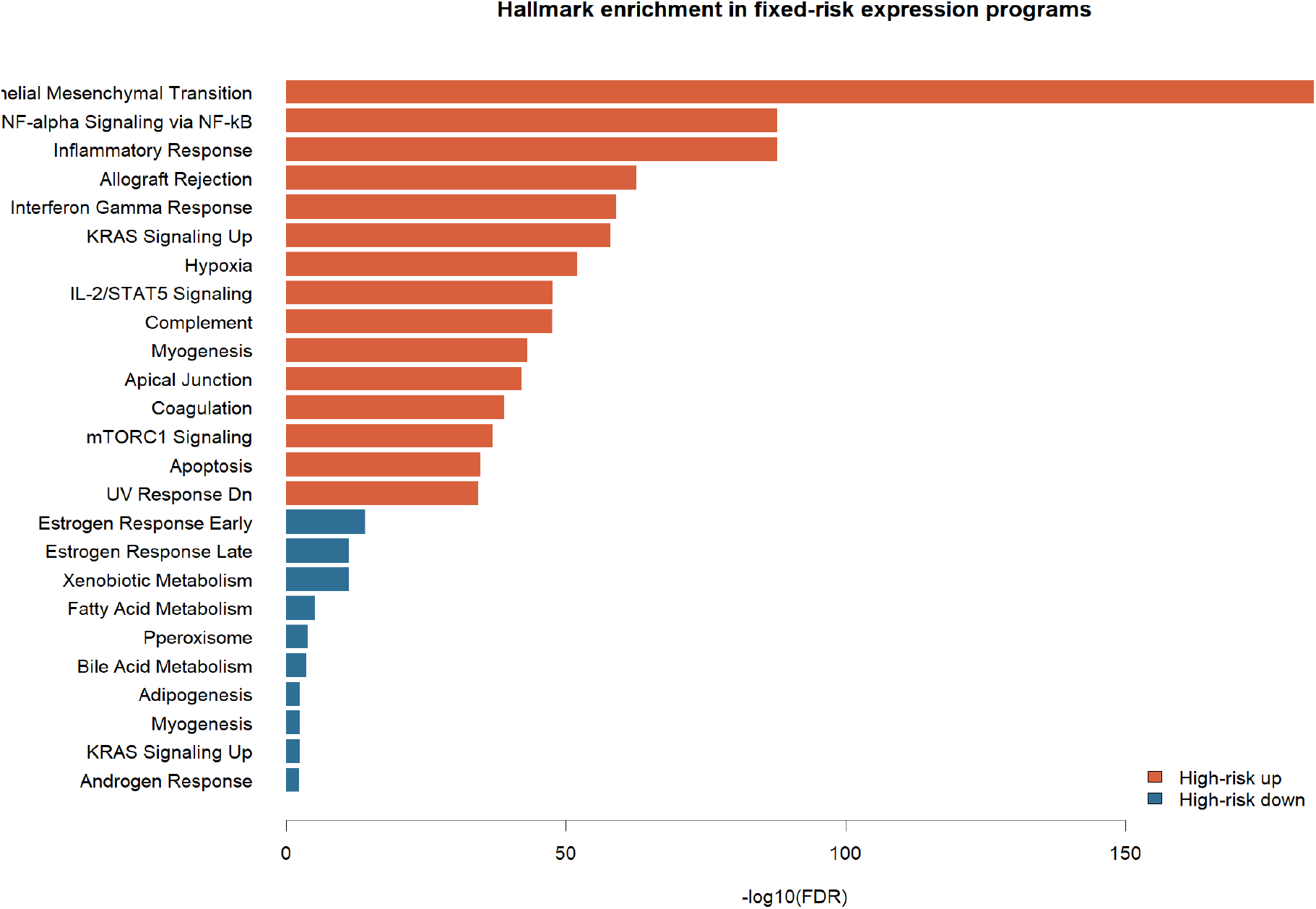
Biological pathway enrichment associated with the fixed risk score.

Supplementary Figures S21-S24 expand the tumor-normal differential expression analysis that underpins the biological interpretation of the score. Supplementary Figure S21 shows the global differential-expression landscape, Supplementary Figure S22 displays the top 50 tumor-normal genes in a heatmap, Supplementary Figure S23 ranks the largest absolute log_2_ fold-change genes, and Supplementary Figure S24 highlights the top tumor-enriched genes. Together, these panels show that the modeling work was evaluated against the broader bladder tumor transcriptional background rather than interpreted in isolation.

Supplementary Figures S25 and S26 extend the risk-group biological analysis after the five-gene score was fixed. Supplementary Figure S25 identifies genes differentially expressed between high-risk and low-risk TCGA tumors, and Supplementary Figure S26 compares immune and stromal signature scores across risk groups. These analyses explain why the risk score is interpreted as a stromal-EMT and immune-suppression phenotype rather than as a purely statistical survival score.

The gene composition of the model supported this interpretation. *EMP1* has been associated with epithelial plasticity, extracellular matrix remodeling, and aggressive phenotypes in multiple cancers. *AHNAK* is a large scaffolding protein implicated in cell architecture, migration, and mesenchymal states. *TNFRSF14* and *CLEC2D* are immune-related genes, and *GSDMB* may reflect epithelial lineage and inflammatory context. The combined score therefore plausibly integrates tumor-cell plasticity and immune microenvironment signals.

### Multi-omics analyses suggested a transcriptional microenvironment phenotype rather than high genomic instability

To test whether high-risk tumors were simply more genomically unstable, we compared mutation and CNV burden by risk group (**Figure 3**). Contrary to that expectation, high-risk tumors had lower nonsilent mutation burden than low-risk tumors (*p* = 0.0010). CNV burden was also lower by mean absolute segment amplitude (*p* = 0.000126) and total altered length (*p* = 0.0111). These findings argue against the score being driven primarily by high genomic instability. RPPA analysis identified multiple protein-level differences between high-risk and low-risk tumors, although current RPPA outputs remain indexed by antibody/protein identifiers that require further annotation before mechanistic claims.

**Figure 3.**
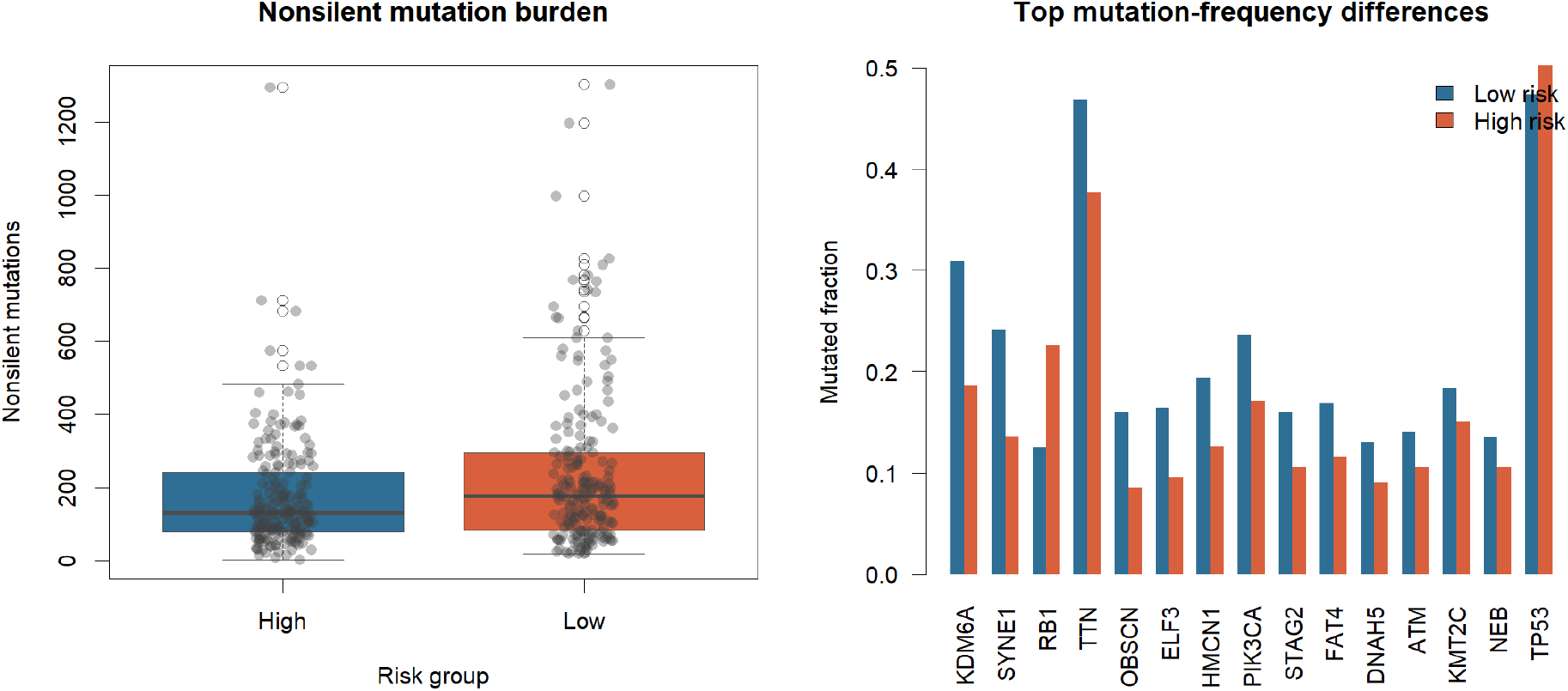
Mutation-level multi-omics support for risk-score-associated tumor biology.

To place the CNV burden difference in a genome-wide context, we also examined the raw distribution of CNV segments across chromosomes in the full TCGA-BLCA cohort (**Figure 4**). Segment counts were highest on chromosome 1 and declined roughly with chromosome size, without a single chromosome showing a disproportionate excess of segments. This genome-wide pattern is consistent with the burden comparison above. The lower CNV burden in high-risk tumors does not reflect a localized hotspot of instability on a particular chromosome, but a broadly lower level of copy-number alteration across the genome, which supports the interpretation that the risk score reflects a transcriptional microenvironment phenotype rather than a chromosome-specific genomic instability signature.

**Figure 4.**
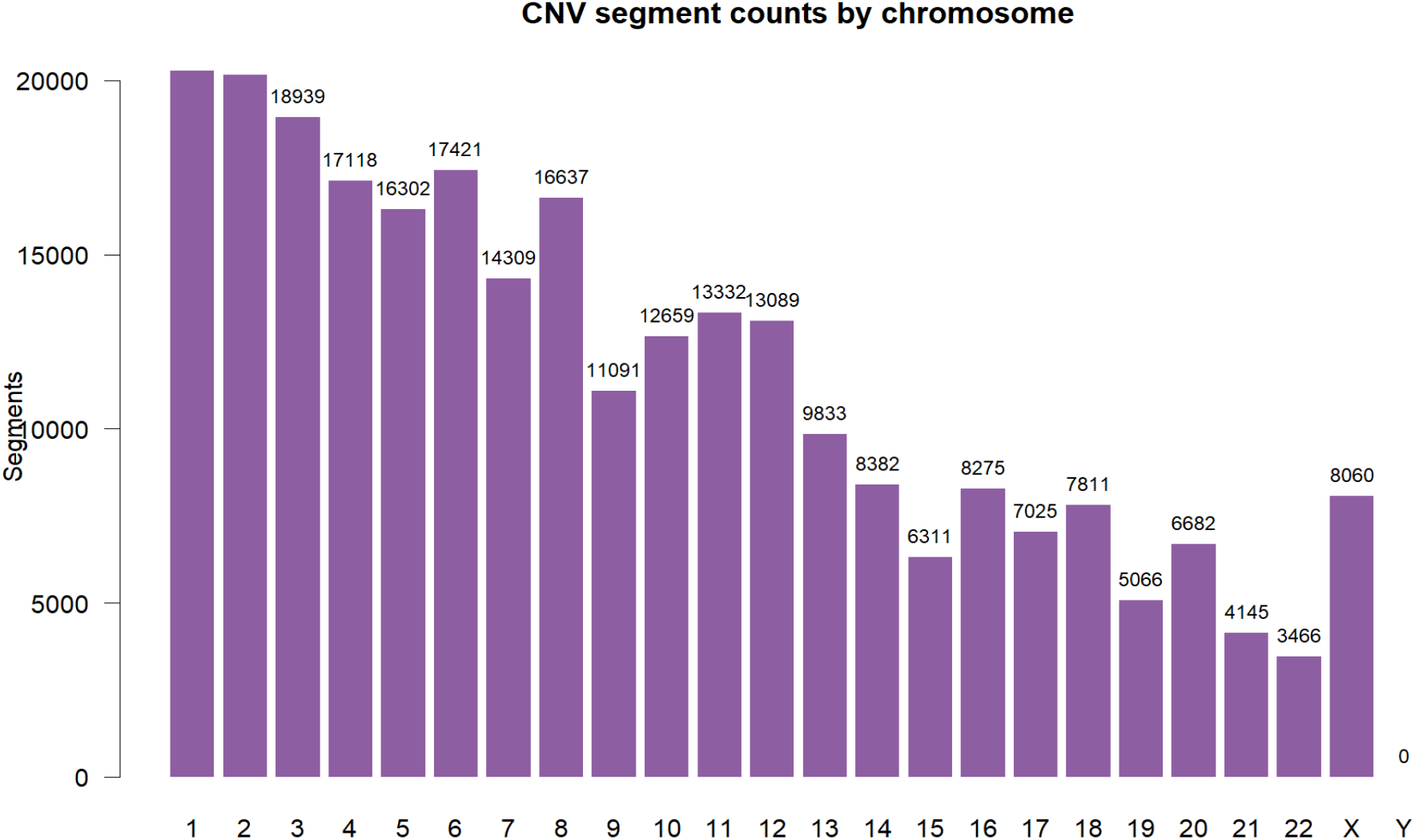
Genome-wide copy-number variation (CNV) segment profile by chromosome across the TCGA-BLCA cohort.

Supplementary Figures S27-S31 provide the expanded multi-omics support for the fixed risk score. Supplementary Figure S27 compares nonsilent mutation burden by risk group, Supplementary Figure S29 examines CNV burden, Supplementary Figure S28 summarizes RPPA protein-feature patterns, Supplementary Figure S30 compares RPPA features between risk groups, and Supplementary Figure S31 displays the most frequently mutated genes in TCGA-BLCA. These panels test whether the score mainly reflects genomic instability or instead captures a transcriptional tumor-microenvironment state.

### Single-cell and spatial datasets localized the score to basal tumor, fibroblast, endothelial, and perivascular compartments

Single-cell analysis provided a cellular explanation for the bulk expression signal (**Figure 5**). In GSE145137, the primary bladder cancer sample contained 14,233 genes and 2,075 cells with author-provided cell-type labels. All five model genes were present. The highest median scores were observed in tumor-labelled urothelial cells, endothelial cells, basal tumor cells, and fibroblasts, whereas T cells had lower scores. Specifically, endothelial cells had median z-score 0.605, basal tumor cells 0.449, fibroblasts 0.337, and T cells -0.418.

**Figure 5.**
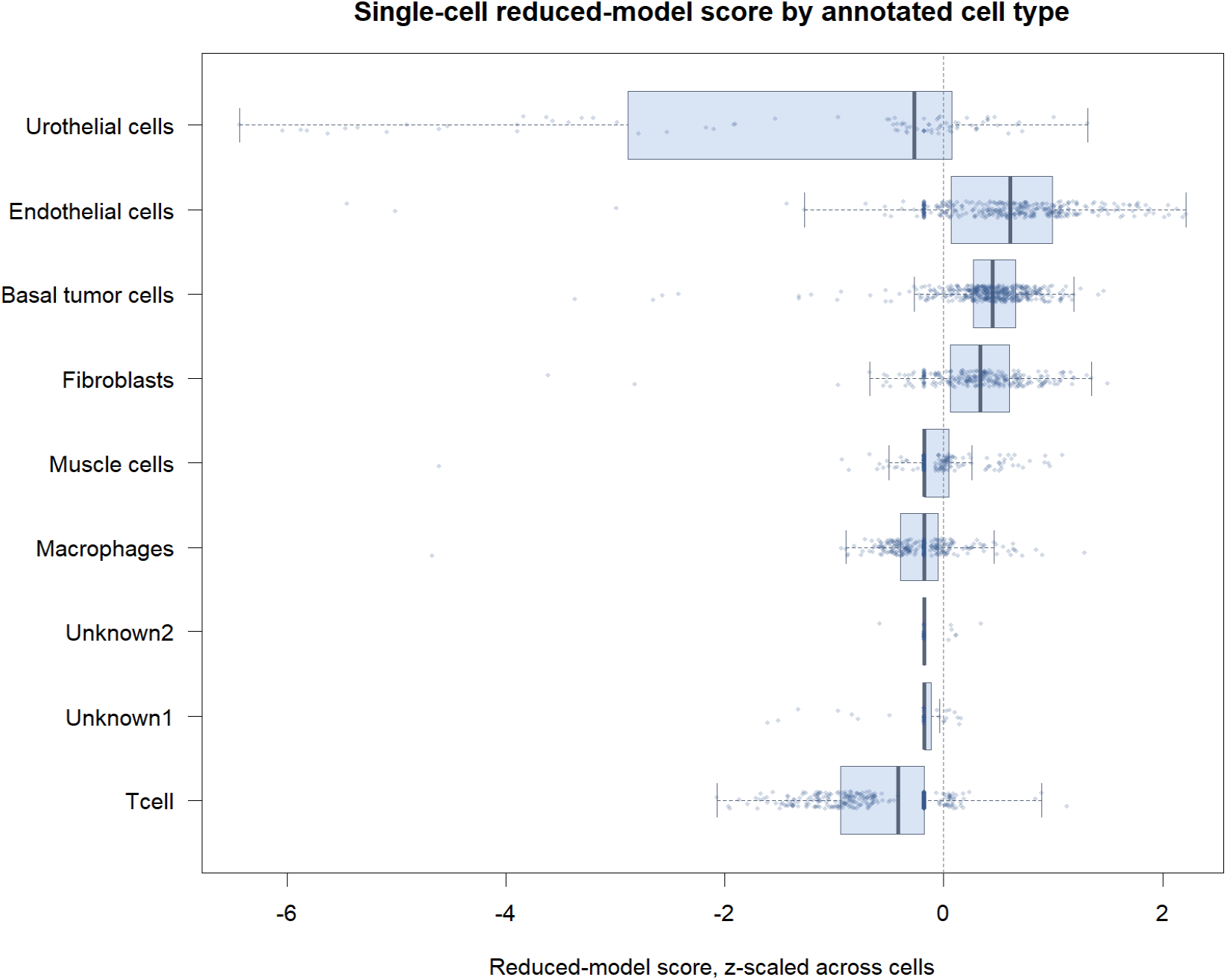
Single-cell localization of the risk score across bladder cancer cell compartments.

The same pattern was reproduced in larger candidate datasets using coarse marker-based cell-type inference. GSE222315 included 102,232 QC-passing single cells across 13 samples. Endothelial cells had median score z = 0.966, fibroblasts 0.287, smooth-muscle/pericyte-like cells 0.198, and T/NK cells -0.167. GSE293189 included 33,829 QC-passing cells across 67 samples and showed an even clearer pattern. Endothelial cells median z = 1.066, fibroblasts 0.639, myeloid cells 0.018, and T/NK cells -0.395. Gene-level inspection across these two candidate cohorts showcased that the endothelial and fibroblast enrichment was driven predominantly by *AHNAK* and *EMP1*, whereas *CLEC2D* was comparatively enriched in T/NK cells, indicating that individual model genes track distinct compartments rather than functioning as a single uniform stromal marker (**Figure 6**). At the sample level, per-sample median risk scores in both candidate cohorts were consistently higher in tumor-annotated samples than in normal, adjacent-normal, or unknown-condition samples, with the most pronounced separation in GSE222315 (**Figure 7**). In GSE171351 Visium spatial transcriptomics, basal epithelial spots had median z = 0.701. These findings suggest that the bulk high-risk phenotype arises from basal tumor, stromal, vascular, and perivascular compartments, with relatively lower contribution from T/NK cells.

**Figure 6.**
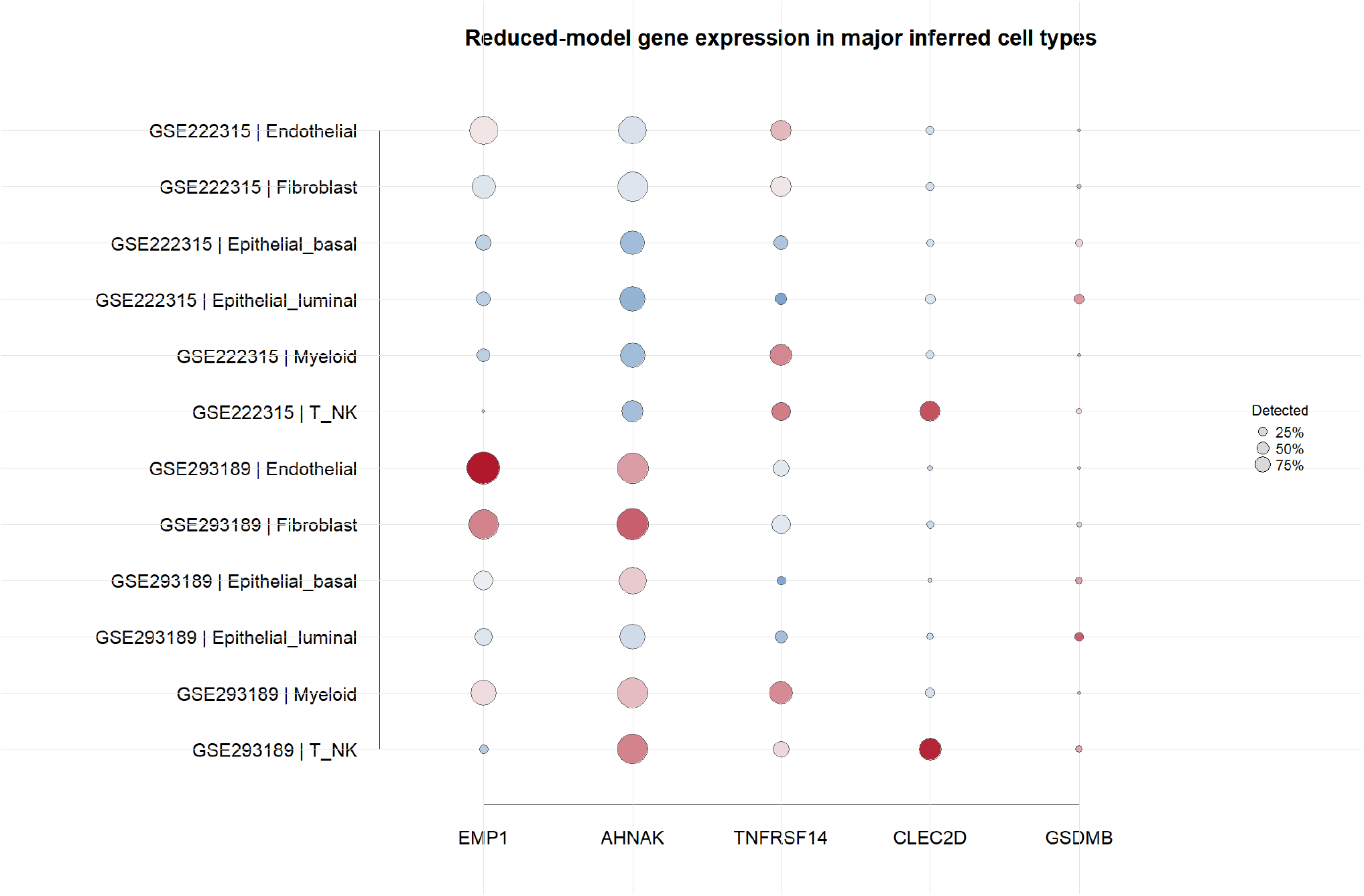
Expression of the five fixed-model genes across annotated cell types in independent candidate single-cell datasets (GSE222315 and GSE293189).

**Figure 7.**
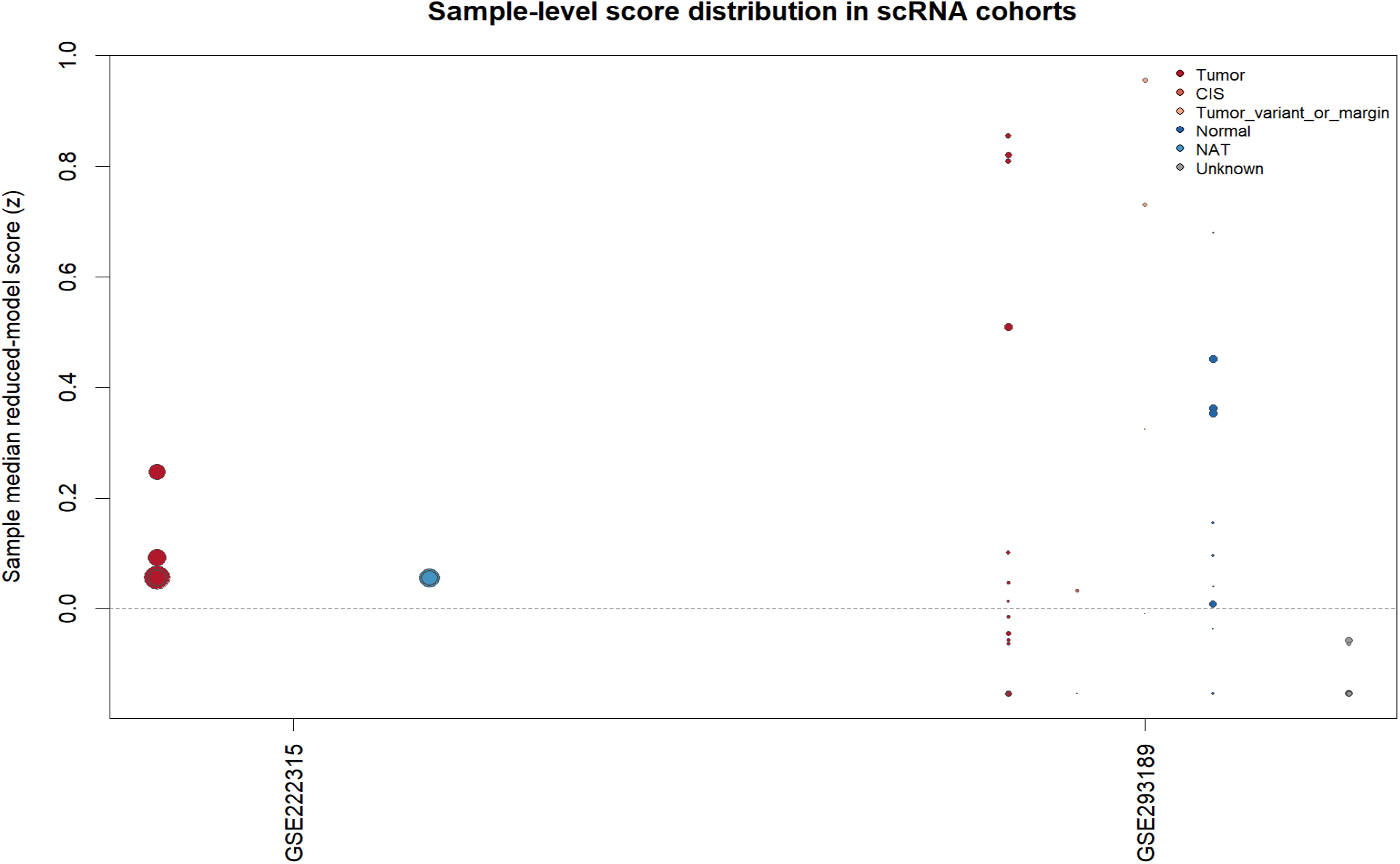
Sample-level distribution of the fixed five-gene risk score by tissue condition in the candidate single-cell/spatial validation cohorts.

Supplementary Figures S32 -S38 provide the single-cell and spatial support for cellular localization of the risk score. Supplementary Figures S32 and S33 summarize cell-type-resolved risk-score patterns in GSE222315 and GSE293189, Supplementary Figure S34 reports the GSE145137 cell-type composition, Supplementary Figure S35 decomposes gene-level risk-score contributions by cell type, Supplementary Figure S36 shows model-gene expression across annotated cells, Supplementary Figure S37 displays the cell-type embedding, and Supplementary Figure S38 maps the fixed risk score onto the single-cell embedding. These analyses show that the bulk score localizes mainly to basal tumor, endothelial, fibroblast, and perivascular compartments, supporting a microenvironment-based interpretation.

### The five-gene score was associated with ICB non-response and poorer OS in IMvigor210

Because the score captured stromal/EMT immune biology, we next tested whether it predicted ICB response. IMvigor210 contained 348 atezolizumab-treated urothelial carcinoma samples, including 298 response-evaluable cases. All five model genes were present. Risk scores were significantly higher in non-responders than responders (Wilcoxon *p* = 0.044), and the AUC for risk score predicting non-response was 0.580. Median-split high-risk tumors had a lower responder rate than low-risk tumors (17.6% versus 28.0%).

Logistic regression supported this association. In IMvigor210, the univariate risk-score odds ratio for non-response was 7.64 (*p* = 0.029). In a multivariate model adjusted for TMB, ECOG, sex, and immune phenotype, the risk-score odds ratio for non-response was 13.54 (*p* = 0.046). The score was also associated with OS. High-risk patients had significantly worse OS than low-risk patients (log-rank *p* = 0.016). As a continuous variable, risk score had univariate Cox HR = 3.23 (*p* = 0.007). In a multivariate OS model adjusted for TMB, ECOG, sex, and immune phenotype, risk score remained significant (HR = 3.15, *p* = 0.044). These findings identify IMvigor210 as the strongest external validation setting for the model, not because it proves broad prognostic utility, but because it connects the stromal-EMT phenotype to ICB resistance.

### GSE176307 provided directionally consistent but statistically weaker ICB support

GSE176307 included 90 metastatic urothelial carcinoma samples treated with ICB, with response, OS, PFS, *FGFR3* alteration status, TMB, ECOG, gender, and treatment annotations. Of 89 response-evaluable cases, 16 were responders and 73 were non-responders. All five model genes were present. The direction of effect was consistent with IMvigor210. Non-responders had higher median risk score than responders, and the AUC for non-response prediction was 0.576. Nevertheless, the Wilcoxon *p* value was 0.342, and the difference was not statistically significant. The high-risk responder rate was 15.6% compared with 20.5% in low-risk tumors. Survival validation in GSE176307 was also non-significant, with OS C-index 0.462, PFS C-index 0.489, and median-split OS log-rank *p* = 0.647. GSE176307 therefore should be treated as supportive external evidence showing consistent response direction rather than as an independent confirmatory survival cohort.

### Risk score correlated with TGF-beta/CAF and EMT-stromal programs in ICB cohorts

To understand why the risk score was associated with ICB non-response, we scored immune and stromal marker programs in IMvigor210 and GSE176307. In both cohorts, risk score correlated most strongly with TGF-beta/CAF and EMT-stromal signatures. In IMvigor210, Spearman correlations were 0.47 for TGF-beta/CAF and 0.55 for EMT-stromal. In GSE176307, corresponding correlations were 0.41 and 0.46. Risk score correlated negatively with luminal epithelial signature in both cohorts. CD8 effector and checkpoint correlations were weak compared with stromal and EMT programs. To summarize these relationships across both ICB cohorts simultaneously, we generated a circos-style association plot linking cohort, risk group, immune phenotype or *FGFR3* status, CD8/TGF-beta signature balance, and response (**Figure 8**), and a Sankey flow diagram tracing patients from cohort and risk group through immune phenotype or *FGFR3* status to response outcome (**Figure 9**). Both integrative visualizations converged on the same picture that high-risk, TGF-beta/CAF-high, CD8/TGF-beta-low patients disproportionately followed the non-responder trajectory in both IMvigor210 and GSE176307. It reinforced that the risk score’s association with ICB resistance reflected a coherent stromal-immune axis rather than a cohort-specific artifact.

**Figure 8.**
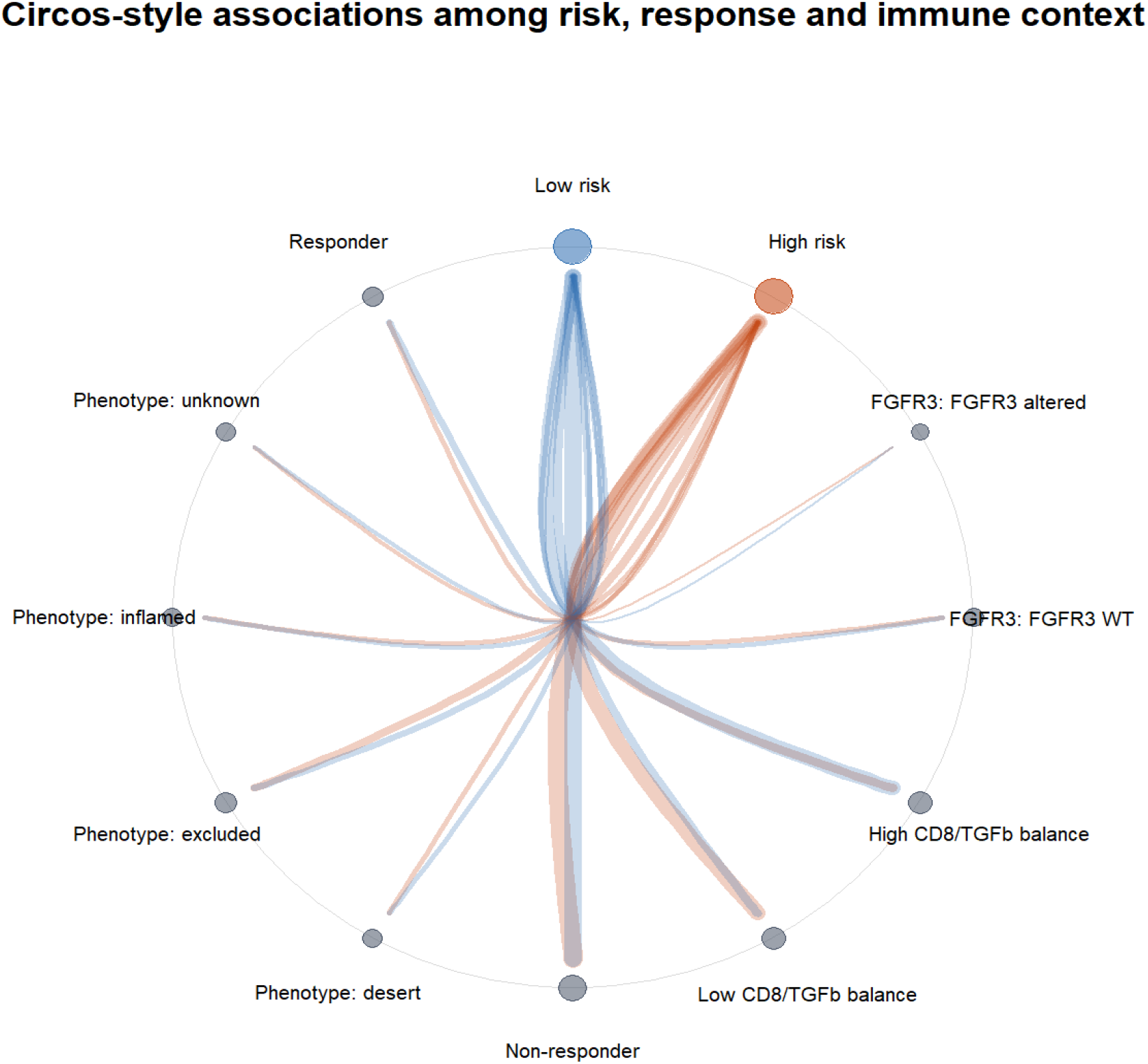
Circos-style association plot summarizing relationships among cohort, risk group, immune phenotype or FGFR3 status, CD8/TGF-beta signature balance, and immunotherapy response in the IMvigor210 and GSE176307 cohorts.

**Figure 9.**
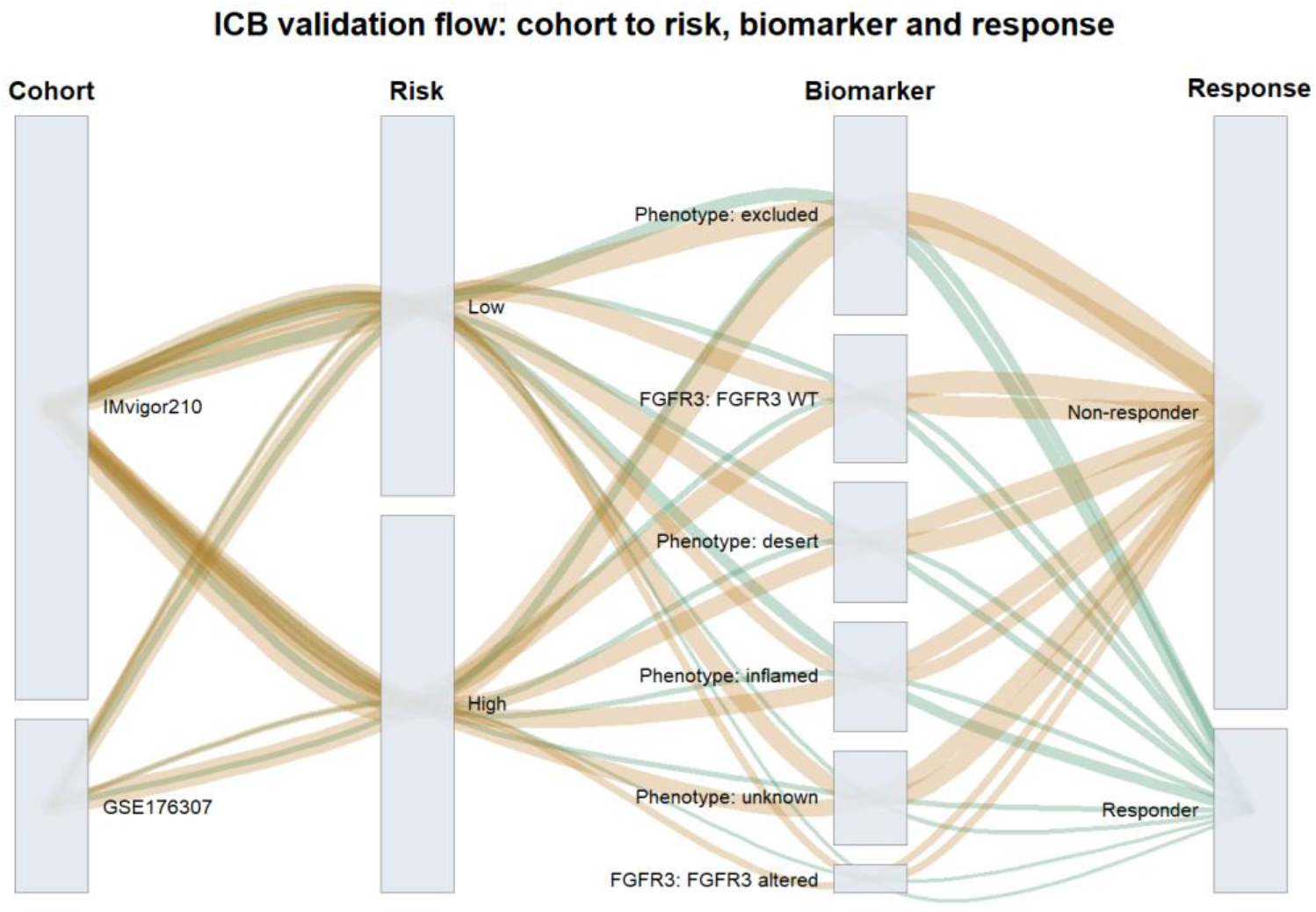
Sankey flow diagram tracing patients from cohort and risk group through immune phenotype or FGFR3 status to immunotherapy response outcome in IMvigor210 and GSE176307.

Supplementary Figures S39-S46 provide the detailed immune-checkpoint-blockade validation layer. Supplementary Figure S39 shows ROC performance for non-response prediction, Supplementary Figure S40 illustrates the cytotoxic/CD8 versus TGF-beta/CAF balance, Supplementary Figures S41 and S42 show OS and PFS survival curves, Supplementary Figure S43 displays model-gene expression patterns across ICB samples, Supplementary Figure S44 compares risk scores between responders and non-responders, Supplementary Figure S45 summarizes response rates by risk group, and Supplementary Figure S46 correlates the risk score with immune, stromal, EMT, and luminal signatures. These panels explain the clinical meaning of the score as an ICB-resistance-associated stromal-EMT program rather than a generic inflammation marker.

### Evidence integration favored a microenvironment-response model over a classical prognostic-signature model

The integrated results illustrated that if the analysis were judged only as a prognostic signature study, the modest GSE13507 and GSE31684 performance would be a serious limitation. However, the same modest prognostic behavior becomes informative when placed beside the pathway, single-cell, and ICB results. The score was internally prognostic in TCGA, externally portable across platforms, strongly associated with stromal/EMT biology, localized to stromal and vascular compartments in single-cell datasets, and associated with ICB non-response in IMvigor210. These layers point to a molecular phenotype that is clinically relevant but not reducible to survival prediction alone.

This interpretation is also consistent with bladder cancer biology. Basal and stromal-rich tumors often have aggressive clinical behavior, but their relationship with therapy is complex. Some immune-infiltrated tumors respond to checkpoint blockade, whereas others remain resistant because T cells are spatially excluded, dysfunctional, or counterbalanced by fibroblast, myeloid, endothelial, or TGF-beta programs. The present score appears to sit in this second category. It does not simply measure CD8 infiltration, nor does it behave as a pure tumor mutation burden surrogate. Instead, it captures an immune-present, stromal/EMT-high state that may blunt effective checkpoint response. This provides a coherent explanation for why the score performed better in IMvigor210 ICB response and OS analysis than in older heterogeneous GEO survival cohorts.

### Relationship to ADC and contemporary treatment hotspots

Current therapeutic development in urothelial carcinoma increasingly involves ICB and ADC combinations. Enfortumab vedotin plus pembrolizumab is now a major reference point for advanced disease, and other ADC strategies continue to evolve. The present study does not include an ADC-treated transcriptomic validation cohort, so ADC response cannot be claimed. Nevertheless, ADC biology is relevant to the framing of the manuscript. ADC activity may depend on target-antigen expression, drug internalization, payload sensitivity, bystander effects, and immune remodeling after tumor-cell injury. A stromal/EMT immune-suppression score could plausibly influence ADC plus ICB outcomes by affecting immune reactivation after ADC-mediated antigen release or immunogenic cell death. In **Figure 10**, we provide drug sensitivity analysis nominating therapeutic vulnerabilities, which are associated with the risk score. Future ADC-treated cohorts should test target genes such as *NECTIN4*, *ERBB2*, and *TACSTD2* together with the five-gene risk score. This preserves relevance to the current ADC hotspot while avoiding a claim that the current dataset cannot support.

**Figure 10.**
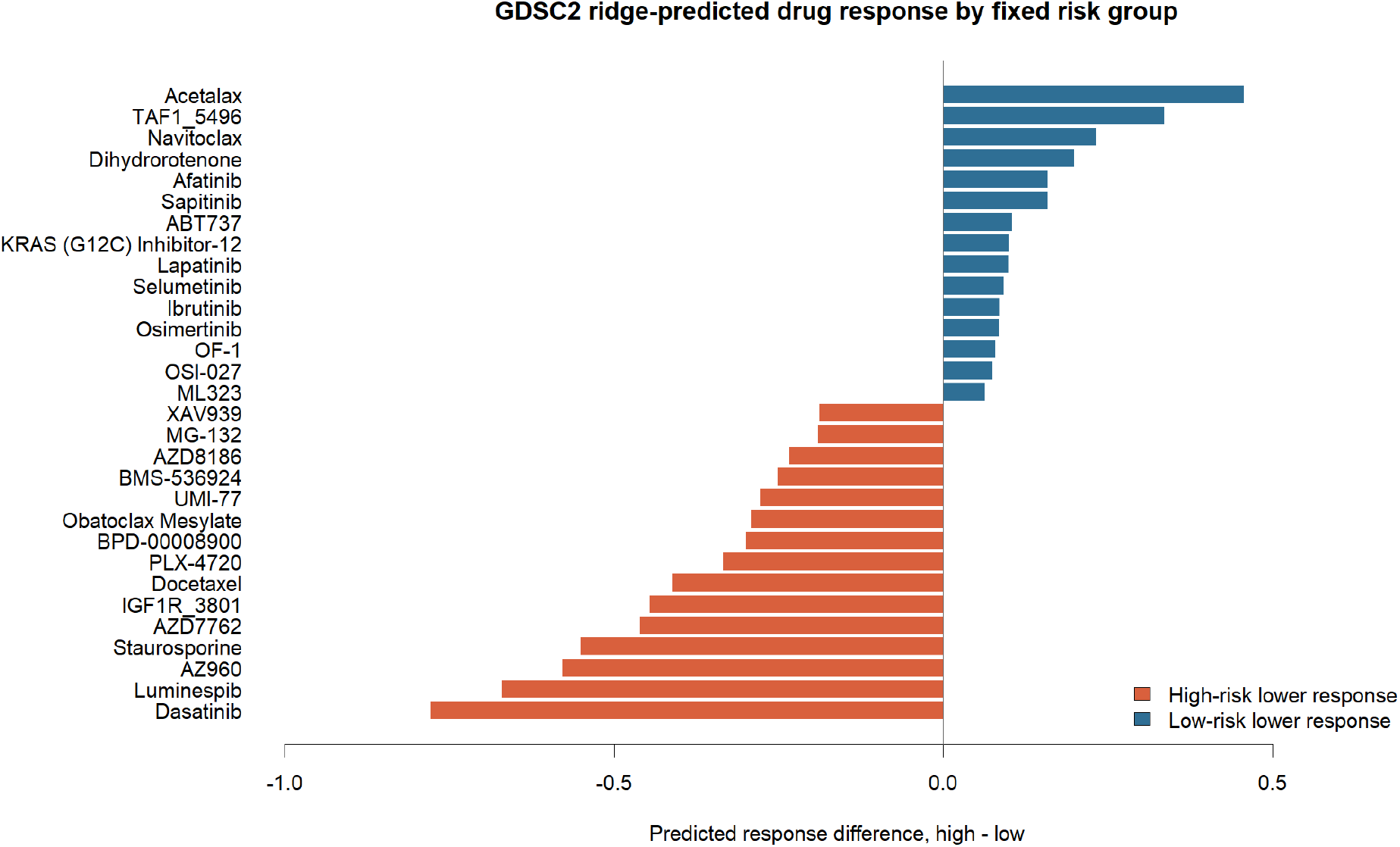
Drug sensitivity analysis nominating therapeutic vulnerabilities associated with the risk score.

### Drug sensitivity analysis nominated high-risk therapeutic hypotheses but did not directly model ADC response

Drug sensitivity prediction using simplified GDSC2 ridge modeling suggested that high-risk tumors may be more sensitive to several agents, including Luminespib, Staurosporine, Obatoclax Mesylate, MG-132, AZ960, Dasatinib, Docetaxel, and Bortezomib (**Figure 10**). These findings are hypothesis-generating and require rerunning with a full oncoPredict environment and experimental validation. Nevertheless, the top-ranked agents overlap with pathways relevant to proteostasis, apoptosis, kinases, cytoskeletal state, and stress signaling, which may plausibly interact with EMT/stromal biology. We did not claim that the five-gene score predicts ADC efficacy. ADC activity depends on target antigen expression, linker-payload biology, internalization, payload resistance, and immune context. However, ADCs are directly relevant to this analysis because enfortumab vedotin plus pembrolizumab is now a central regimen in advanced urothelial carcinoma [14]. The score’s relationship with ICB resistance and stromal immune suppression provides a rationale for future analyses testing whether stromal-EMT states influence ADC plus ICB response, especially in datasets with Nectin-4, HER2, or Trop-2 expression and ADC-treated outcomes.

Because no supplementary ADC-response figure is included, the drug and ADC discussion should be read alongside the main therapeutic-hypothesis figure rather than as direct ADC validation. The supplementary figure set instead strengthens the upstream evidence chain: cohort quality control, differential-expression context, model construction, external validation, multi-omics interpretation, single-cell localization, and ICB-response biology. This structure keeps the therapeutic claim hypothesis-generating while grounding it in the validated transcriptomic and immune-microenvironment analyses.

## Discussion

This study developed and characterized a compact five-gene bladder cancer risk score that connects TCGA prognosis, tumor microenvironment biology, single-cell localization, ICB response, and therapeutic hypotheses. Though the score achieved internal TCGA survival performance, its external survival validation in GSE13507 and GSE31684 was modest. Rather than overstating the score as a definitive prognostic classifier, we reframed it as a transcriptomic readout of stromal-EMT and immune-suppressive biology. Under this framing, the downstream results are coherent and clinically relevant. The five genes in the model plausibly capture multiple compartments. *EMP1* and *AHNAK* are associated with epithelial plasticity, mesenchymal phenotypes, and cell-structure remodeling. *TNFRSF14* and *CLEC2D* point toward immune-interaction biology. *GSDMB* may reflect epithelial lineage and inflammatory context. The model is therefore not merely a statistical artifact. It condenses tumor-cell and microenvironment signals into a portable score. The single-cell results strengthen this interpretation by showing that the score is highest in basal tumor, endothelial, fibroblast, and smooth-muscle/pericyte-like compartments, and lower in T/NK cells. Extending this localization to two additional, independently processed single-cell cohorts using marker-based cell-type inference reproduced the same compartment pattern, which showed that the enrichment is carried mainly by *AHNAK* and *EMP1*, while *CLEC2D* instead marks the T/NK compartment. This gene-level and sample-level consistency across three separate single-cell datasets argues against a batch- or cohort-specific artifact and supports a reproducible cellular basis for the bulk risk score.

It is also important to situate this five-gene signature within the broader landscape of previously published bladder cancer prognostic gene-expression models, several of which have used similar TCGA/GEO-based Cox-regression frameworks to nominate small gene panels [35, 36]. Notably, three of the five genes retained in our model, *GSDMB*, *CLEC2D*, and *TNFRSF14*, were previously reported as components of an independent five-gene prognostic signature (together with *APOL2* and *GBP2*) derived from TCGA, GSE13507, GSE32894, and the Mariathasan IMvigor210 cohort, in which the role of *TNFRSF14* was additionally validated experimentally in bladder cancer cell lines [37]. Separately, *EMP1* and *AHNAK*, the two genes carrying positive coefficients in our model, were previously identified as two of six components of a tumor-infiltrating T-cell/neutrophil-derived prognostic signature validated across three independent GEO cohorts and a hospital cohort [38]. Our model therefore does not introduce five entirely novel prognostic genes. Rather, each gene has individually accrued independent prior evidence of prognostic relevance in bladder cancer, and the contribution of the present work lies in (i) identifying this particular five-gene combination as the subset shared across TCGA, GSE13507, and GSE31684 for cross-platform compatibility, and (ii) extending its characterization substantially beyond prior work to include single-cell and spatial compartment localization and, most notably, direct validation against ICB response in IMvigor210 and GSE176307, neither of which was examined in the prior *GSDMB*/*CLEC2D*/*TNFRSF14*-containing or *EMP1*/*AHNAK*-containing signatures. We view the convergence of two independently derived prior signatures on overlapping subsets of our five genes as external, non-circular support for their biological relevance, while acknowledging that the novelty of the present work rests primarily on the immune-checkpoint-response and multi-compartment characterization rather than on the discovery of the individual genes themselves.

The ICB validation is the strongest clinically oriented finding. IMvigor210 showed that high risk was associated with non-response, lower responder rate, and worse OS after atezolizumab. The association remained significant in multivariate logistic and Cox models adjusted for relevant covariates. The effect sizes should be interpreted cautiously because the score is continuous, confidence intervals are wide, and the study remains retrospective. Still, the consistency among response, survival, and stromal/TGF-beta correlations supports a real biological signal. GSE176307 did not reach significance, but the response direction and AUC were similar to IMvigor210, suggesting that the relationship may be present but underpowered in this smaller, heterogeneous real-world cohort. Viewing both cohorts together through the circos-style association plot and the Sankey response-flow diagram makes this pattern easier to grasp at a glance: high-risk, stromal/TGF-beta-skewed patients consistently funnel toward non-response in both datasets, which is a useful way to communicate a multi-cohort, multi-variable resistance signal that would otherwise require several separate statistical statements. These findings align with prior work showing that TGF-beta-associated fibroblast programs can exclude T cells and limit PD-L1 blockade response in urothelial carcinoma [19, 20]. Notably, our high-risk tumors were not simply immune-cold. They showed inflammatory, checkpoint, macrophage, and cytotoxic signatures alongside strong CAF and EMT programs. This distinction matters. Tumors may contain immune infiltrates but remain functionally resistant if stromal architecture, TGF-beta signaling, macrophage polarization, endothelial barriers, or mesenchymal tumor-cell states prevent productive cytotoxic activity. The risk score may therefore capture an immune-present but suppressed phenotype rather than a simple immune desert.

The multi-omics findings further refine the study. High-risk tumors had lower nonsilent mutation burden and lower CNV burden, arguing against the score being a surrogate for genomic instability. The genome-wide CNV segment distribution supports this interpretation further. Segment counts declined smoothly with chromosome size rather than clustering on a particular chromosome, so the lower CNV burden in high-risk tumors reflects a broadly quieter genome rather than a localized instability hotspot that a per-chromosome burden comparison alone might have missed. This may help explain why the score relates to ICB response in a way that is not redundant with TMB. In IMvigor210 multivariate models, TMB was protective, as expected, while the risk score remained associated with non-response and OS. The model may therefore identify a resistance axis orthogonal to mutation-derived neoantigenicity. The ADC context is important but should be handled carefully. Current urothelial carcinoma therapy is increasingly shaped by ADC-based regimens, especially enfortumab vedotin plus pembrolizumab [14]. ADCs may interact with antitumor immunity through immunogenic cell death, antigen release, bystander effects, and microenvironment remodeling. However, our current analysis did not include an ADC-treated validation cohort and should not claim ADC-response prediction. The correct manuscript position is that the score identifies a stromal-EMT immune-resistance state that is relevant to modern ICB and ADC-combination questions. Future ADC datasets should test whether high-risk tumors differ in Nectin-4, HER2, Trop-2, payload sensitivity, or EV plus pembrolizumab benefit.

Drug sensitivity analysis generated additional hypotheses. Several agents predicted to have greater activity in high-risk tumors target stress response, apoptosis, proteasome function, kinases, or cytoskeletal states. Because the local programming environment required a simplified predictor rather than full oncoPredict, these results should be placed as supportive or supplementary unless reproduced in a full pharmacogenomic workflow. Drug analysis is useful for generating mechanistic hypotheses. From a translational perspective, the most realistic near-term use of this score is not immediate clinical decision-making. Rather, it can be used as a compact biological stratifier in retrospective trial datasets and future prospective correlative studies. For example, in ICB cohorts it could be tested alongside PD-L1, TMB, molecular subtype, TGF-beta signatures, and spatial immune-exclusion metrics. In ADC plus ICB cohorts, it could be tested as a modifier of response among tumors with adequate ADC target expression. In single-cell and spatial studies, it can help identify whether stromal, endothelial, pericyte, or basal tumor compartments dominate the resistant phenotype. These applications are more defensible than proposing the score as a standalone clinical assay at this stage.

The five-gene model is fixed and cross-platform compatible. TCGA internal validation is positive, and high risk is consistently stromal/EMT and TGF-beta/CAF associated. Single-cell and spatial analyses localize the score to basal/stromal/vascular compartments. IMvigor210 supports association with ICB non-response and OS.

This study has limitations. First, all analyses are retrospective and based on public datasets with variable preprocessing, treatment, stage distribution, and follow-up. Second, the TCGA-derived model has only modest GEO survival validation, limiting claims about broad prognostic utility. Third, single-cell analyses for some datasets used marker-based coarse labels rather than author-provided annotations, and GSE171351 is spatial spot-level rather than single-cell data. Fourth, RPPA features require improved antibody annotation before detailed protein-level claims. Despite these limitations, the integrated evidence is strong. TCGA internal validation establishes a survival-associated score. GEO validation constrains the claims and encourages caution. Pathway, immune, and single-cell analyses explain the biological state captured by the score. IMvigor210 validates clinical relevance to ICB response. GSE176307 adds directionally consistent external support. Drug prediction and ADC discussion position the work within current therapeutic developments without overclaiming. This combination makes current presentation a multi-omics and immunotherapy-response study rather than a conventional prognostic-signature investigation.

## Conclusion

We developed a fixed five-gene TCGA-derived bladder cancer score composed of *EMP1*, *AHNAK*, *TNFRSF14*, *CLEC2D*, and *GSDMB*. The score showed internal prognostic value in TCGA but only modest external survival validation in GEO. Its strongest value lies in identifying a stromal-EMT and immune-suppressive phenotype that localizes to basal tumor, fibroblast, endothelial, and perivascular compartments and is associated with ICB non-response and worse OS in IMvigor210. These findings support our investigation focused on tumor microenvironment biology and immunotherapy resistance, with ADC and drug-sensitivity analyses framed as clinically relevant future directions.

