## supplementary materials for "A Five-Gene Stromal-EMT Signature Predicts Prognosis, Immunotherapy Resistance, and Therapeutic Vulnerability in Bladder Cancer"

This document contains supplementary figures that support the analyses described in the main text. These include the TCGA-BLCA cohort/data-availability overview, the tumor-normal differential expression landscape, the original exploratory 14-gene TCGA model, the fixed five-gene model coefficient plot, expanded GEO and RPPA detail, the composite IMvigor210/GSE176307 ROC validation figure, and additional single-cell detail for GSE145137.

### Supplementary Figures

*Supplementary Figure S1. TCGA-BLCA cohort and multi-omics data availability used for integrative analysis, summarizing sample counts and data types (RNA-seq, clinical, mutation, CNV, RPPA) available for each analysis module.*

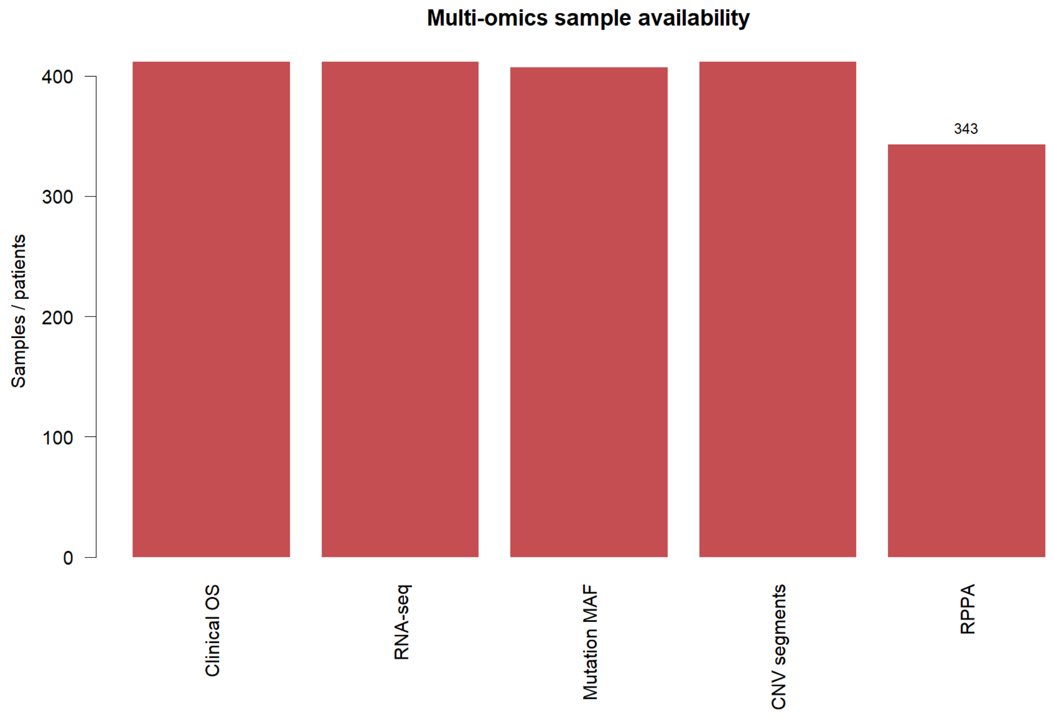

*Supplementary Figure S2. Distribution of AJCC clinical stage among TCGA-BLCA tumor cases used in this study.*

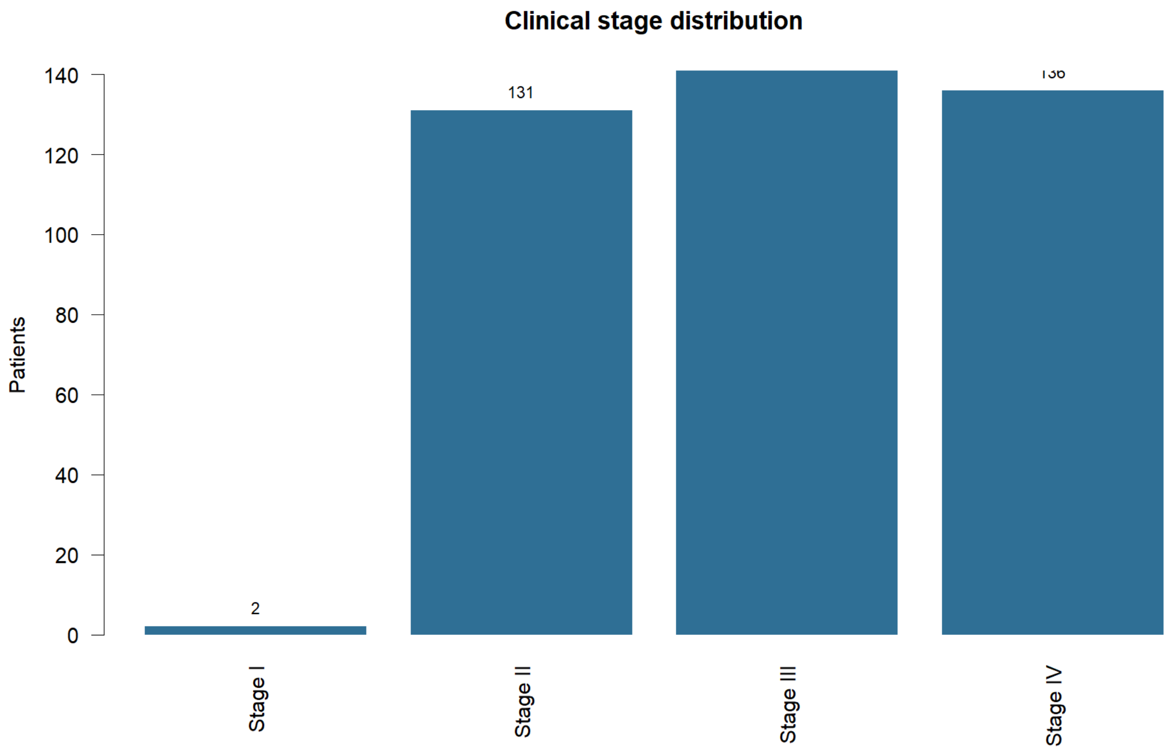

*Supplementary Figure S3. Principal component analysis (PCA) of TCGA-BLCA RNA-seq samples, colored by tumor versus normal tissue type.*

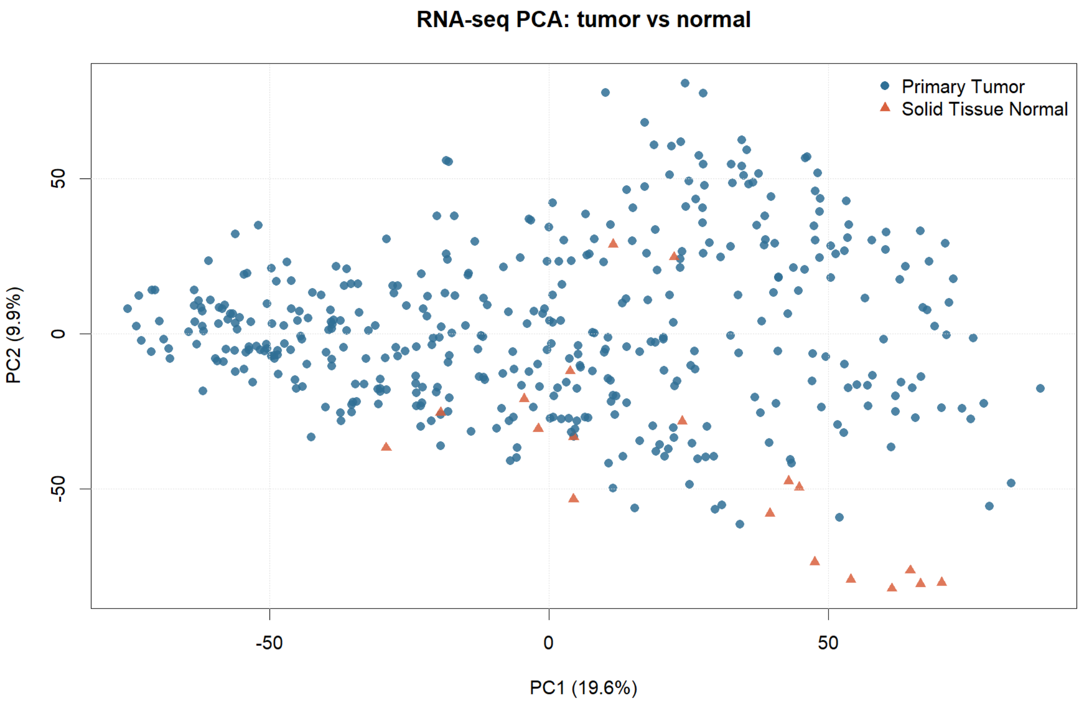


*Supplementary Figure S4. Bar plot of the number of TCGA-BLCA RNA-seq samples by tissue type (tumor versus normal).*

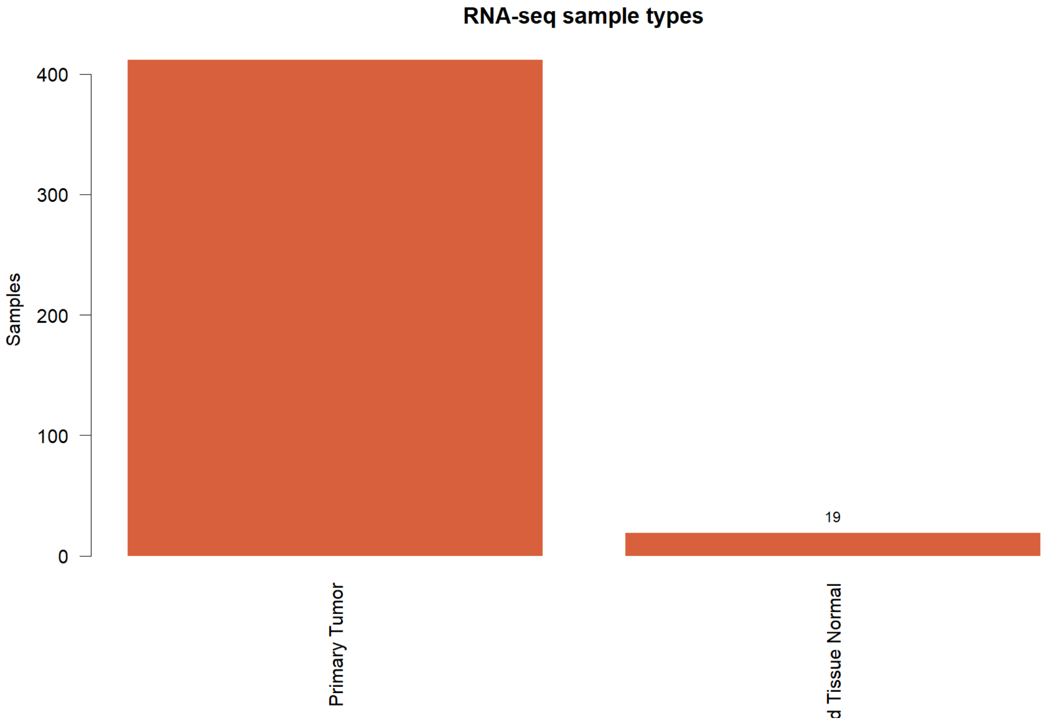

*Supplementary Figure S5. Kaplan-Meier overall survival curves for the exploratory (original 14-gene) TCGA risk score, stratified by median risk group.*

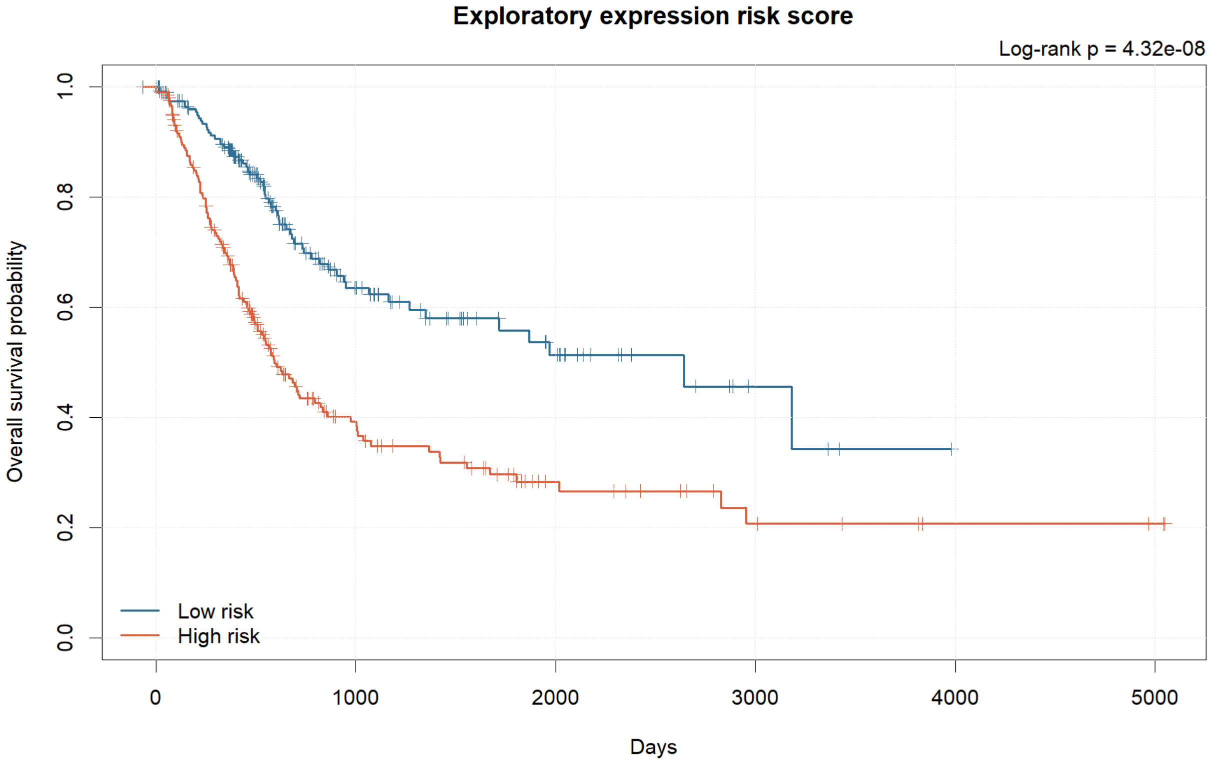

*Supplementary Figure S6. Distribution of the exploratory 14-gene TCGA risk score across tumor samples, shown prior to reduction to the final GEO-compatible five-gene model.*

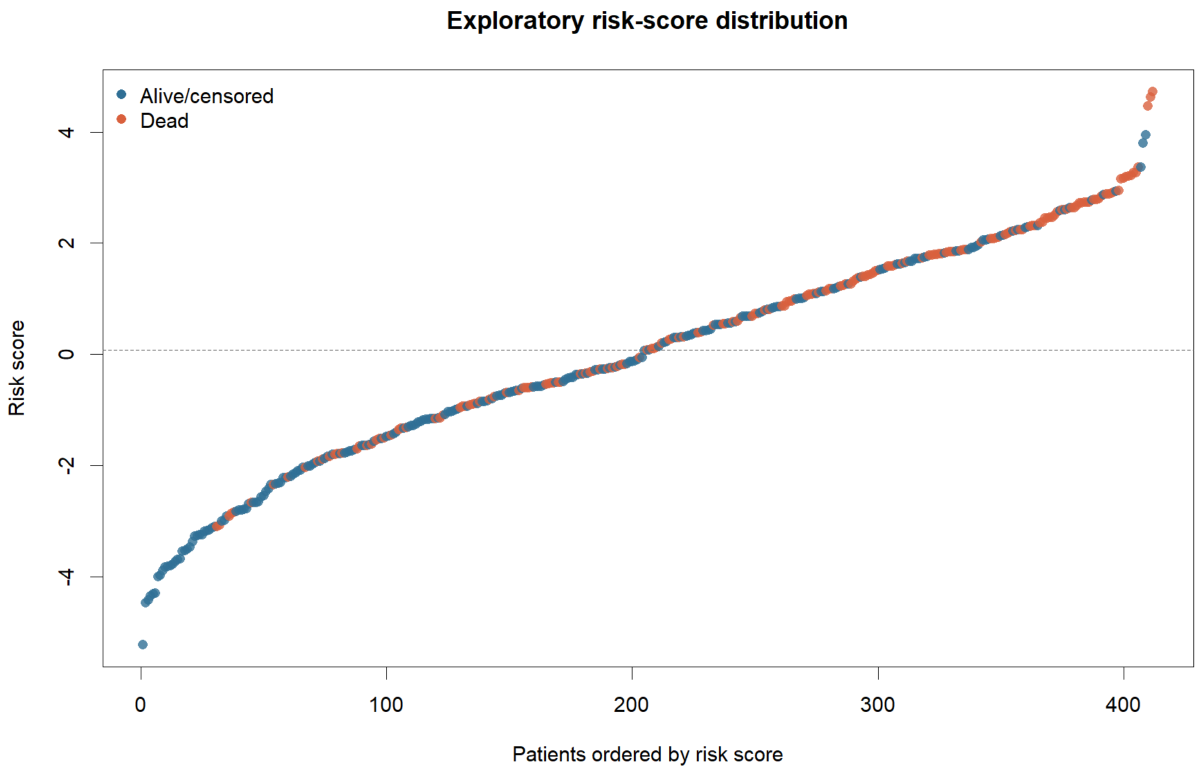

*Supplementary Figure S7. Kaplan-Meier overall survival curves for the top individual prognosis-associated candidate genes identified during univariate screening.*

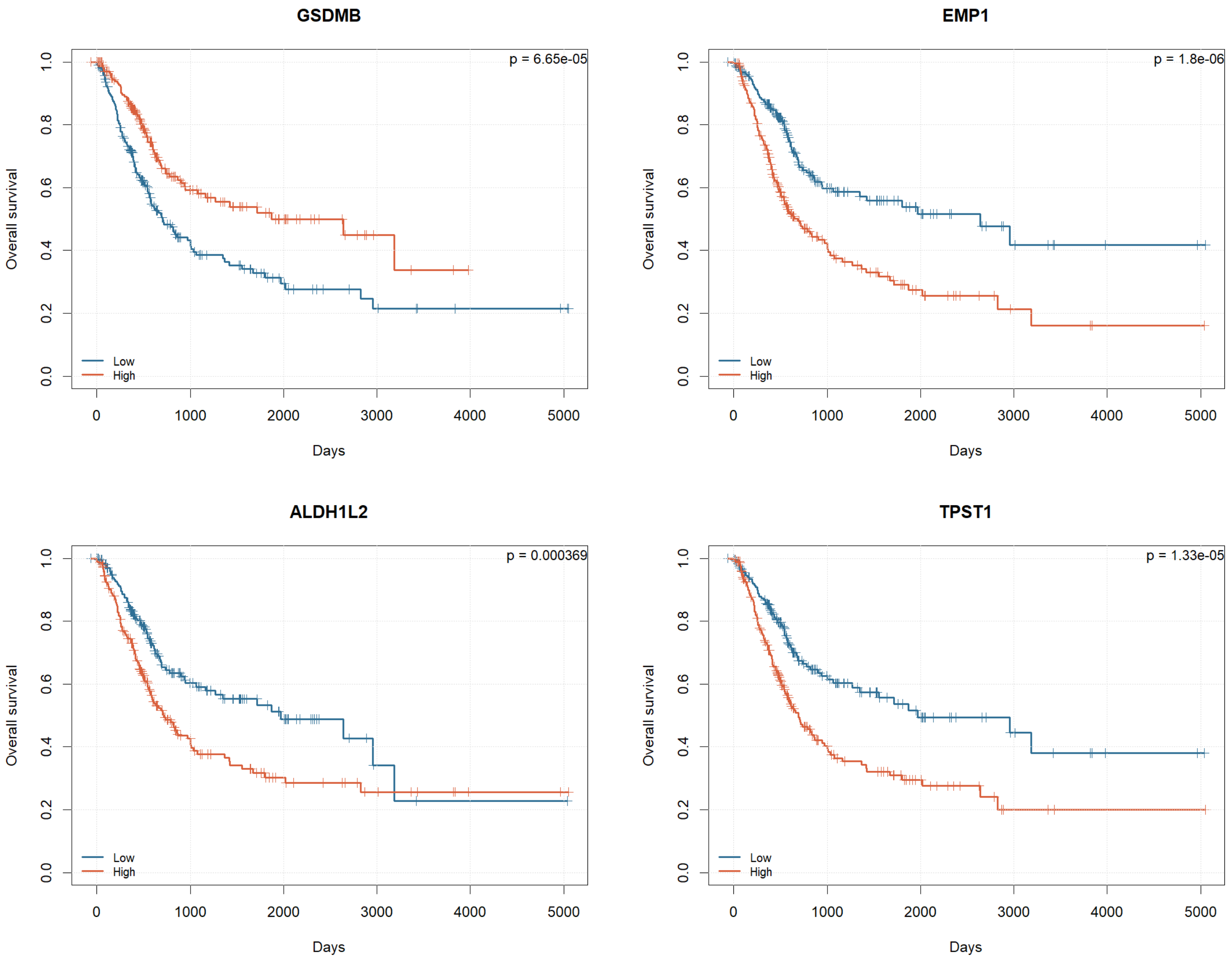

*Supplementary Figure S8. Volcano plot of univariate Cox hazard ratios and significance levels for candidate prognostic genes screened in the TCGA training set.*

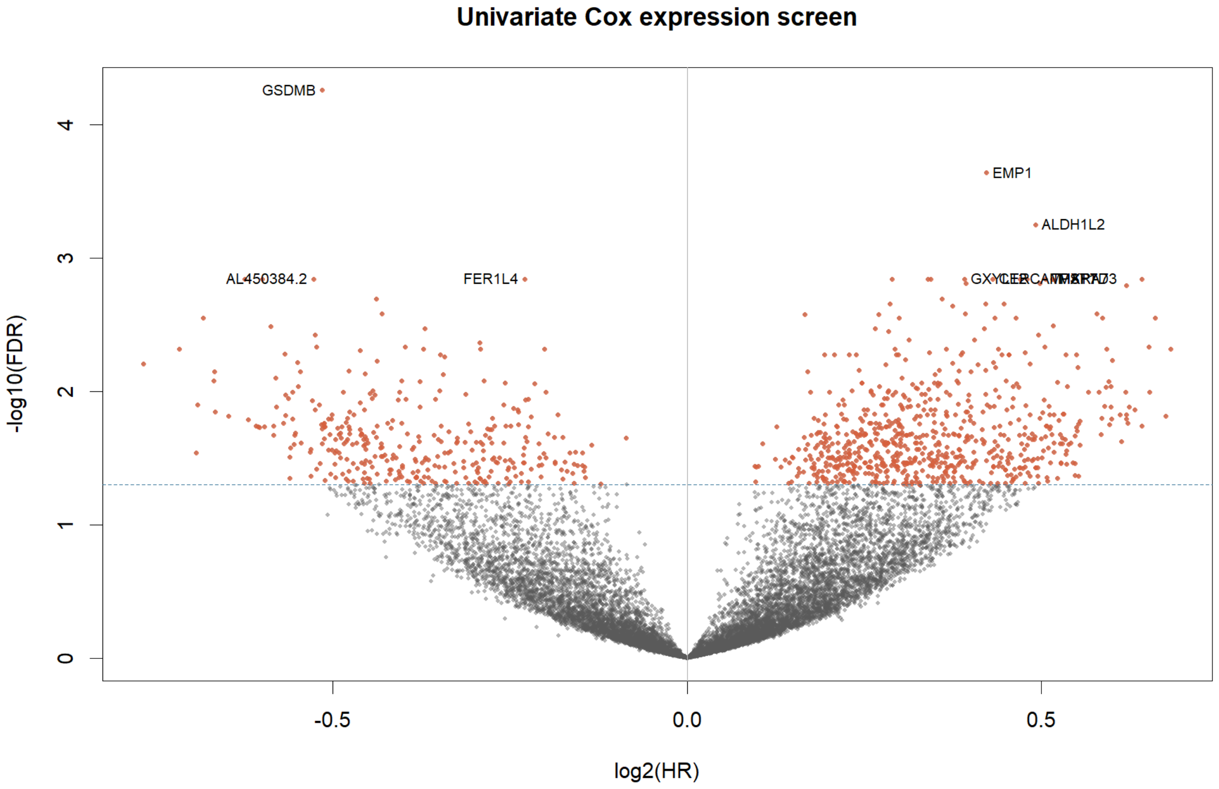

*Supplementary Figure S9. Bar plot of LASSO-Cox coefficients from the original 14-gene TCGA model, prior to reduction to the GEO-compatible five-gene score. (Corrected from the previous version, which mislabeled this panel as the fixed five-gene model.)*

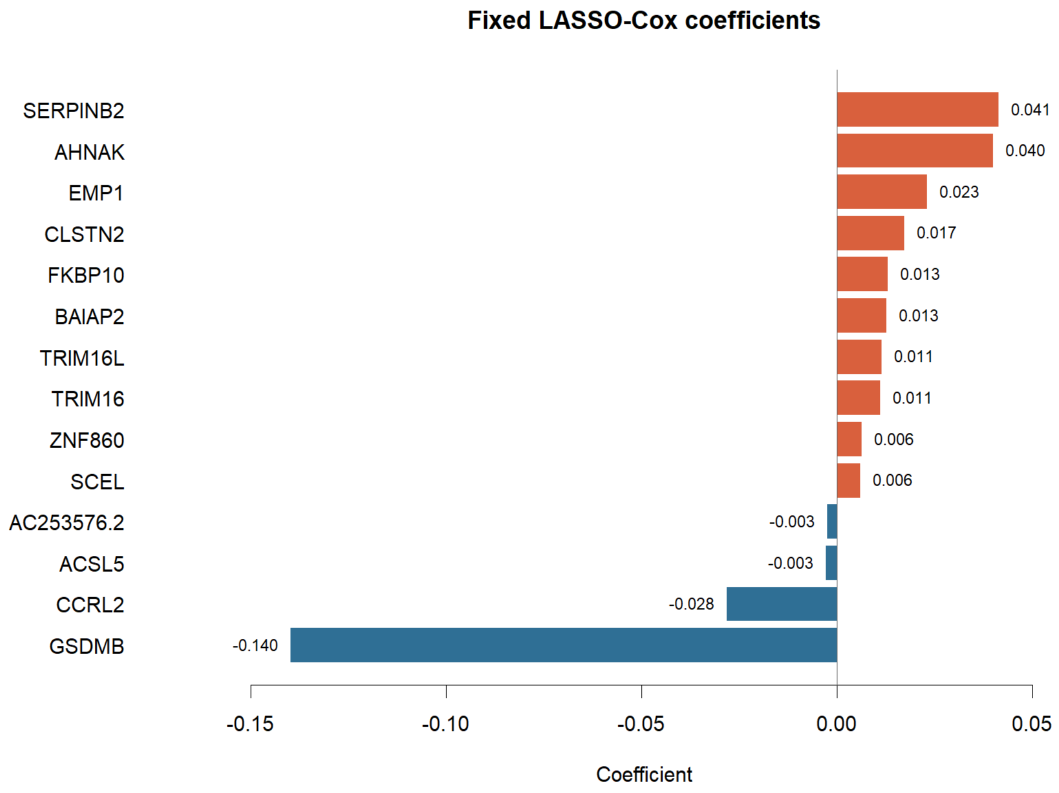

*Supplementary Figure S10. Fixed reduced five-gene model and gene coefficients derived from TCGA, showing the final GEO-compatible five-gene formula after restricting candidate genes to those shared across TCGA, GSE13507, and GSE31684.*

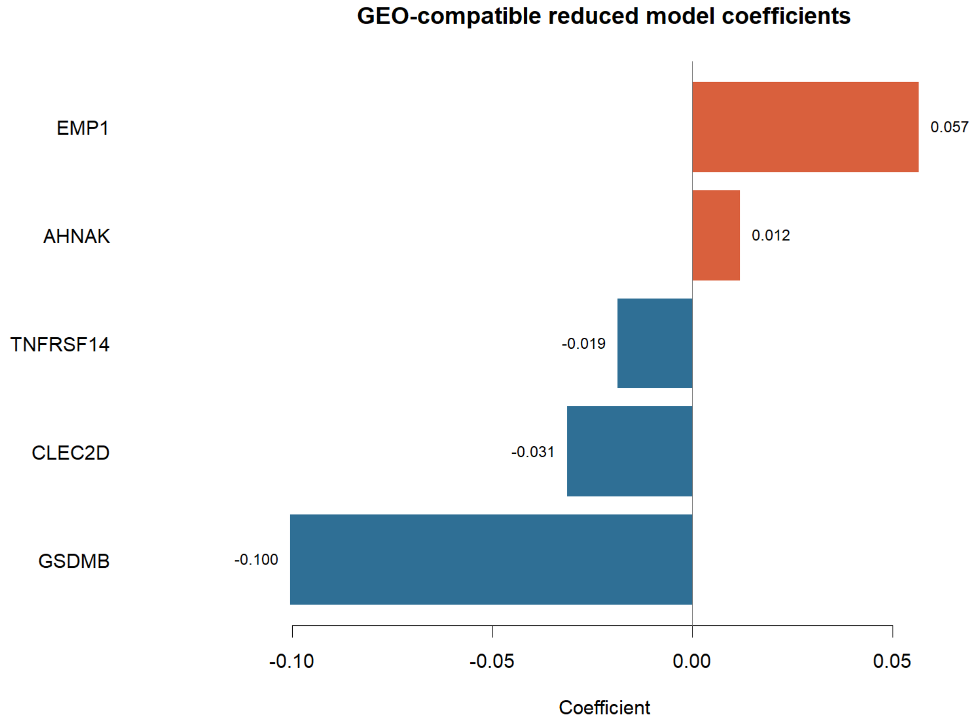


*Supplementary Figure S11. Kaplan-Meier overall survival curves for TCGA-BLCA patients stratified by sex.*

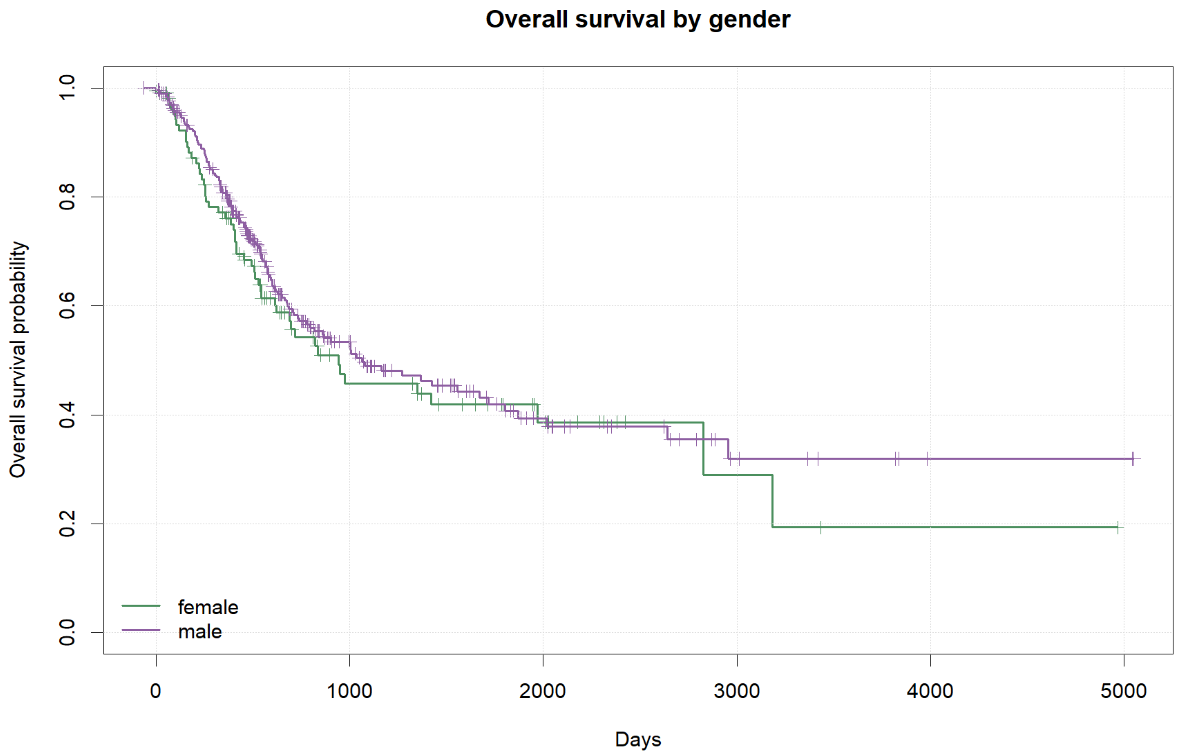
Overall survival by gender
Overall survival probability
0.0
0.2
0.4
0.6
0.8
1.0
0
1000
2000
3000
4000
5000
Days
female
male

*Supplementary Figure S12. Kaplan-Meier overall survival curves for TCGA-BLCA patients stratified by AJCC clinical stage.*

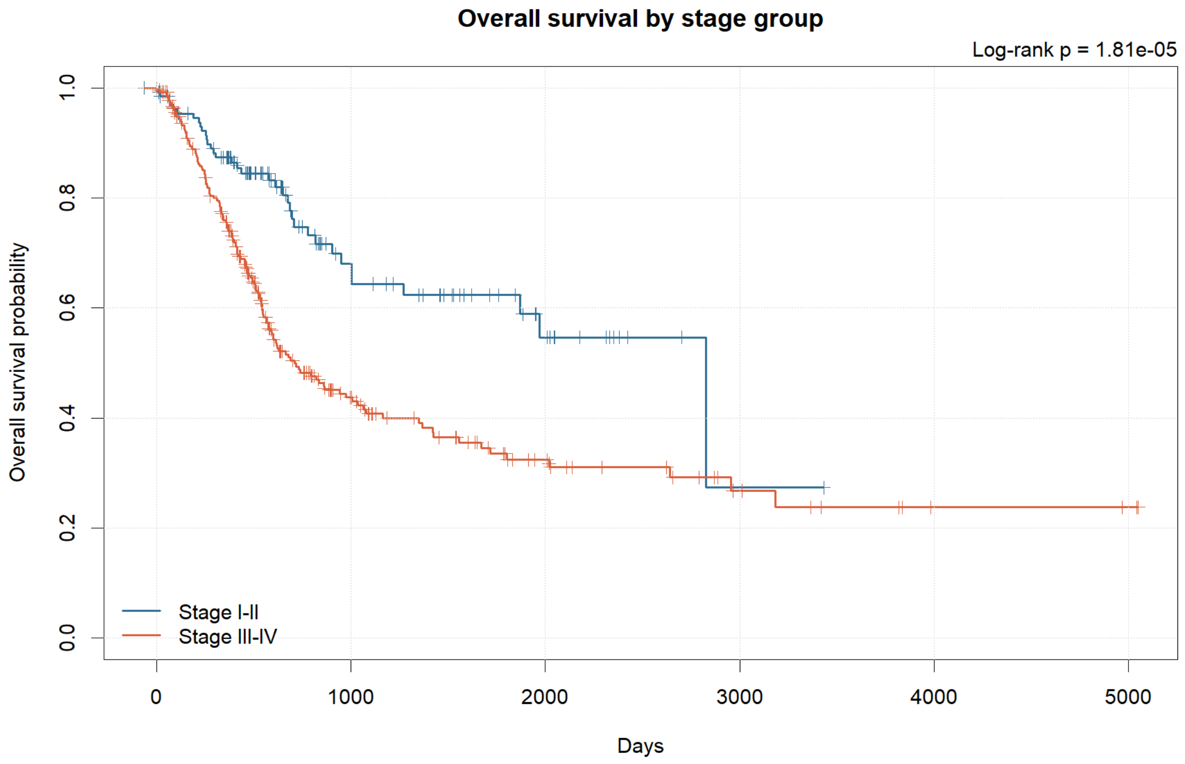

*Supplementary Figure S13. Time-dependent ROC curves for the fixed five-gene model evaluated in the TCGA training, testing, and all-sample sets.*

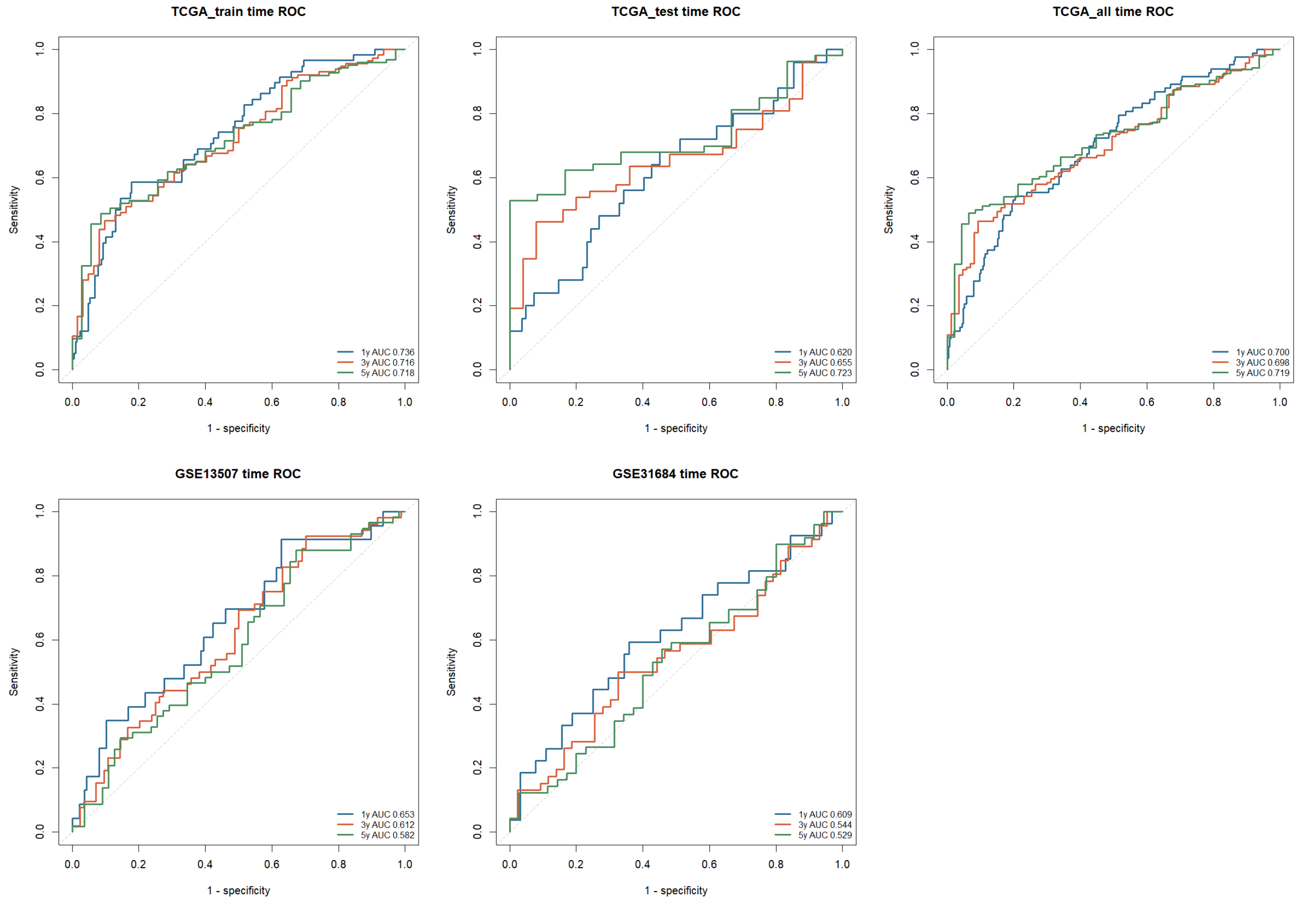

*Supplementary Figure S14. Kaplan-Meier overall survival curves for the fixed five-gene model in the TCGA training, testing, and all-sample sets.*

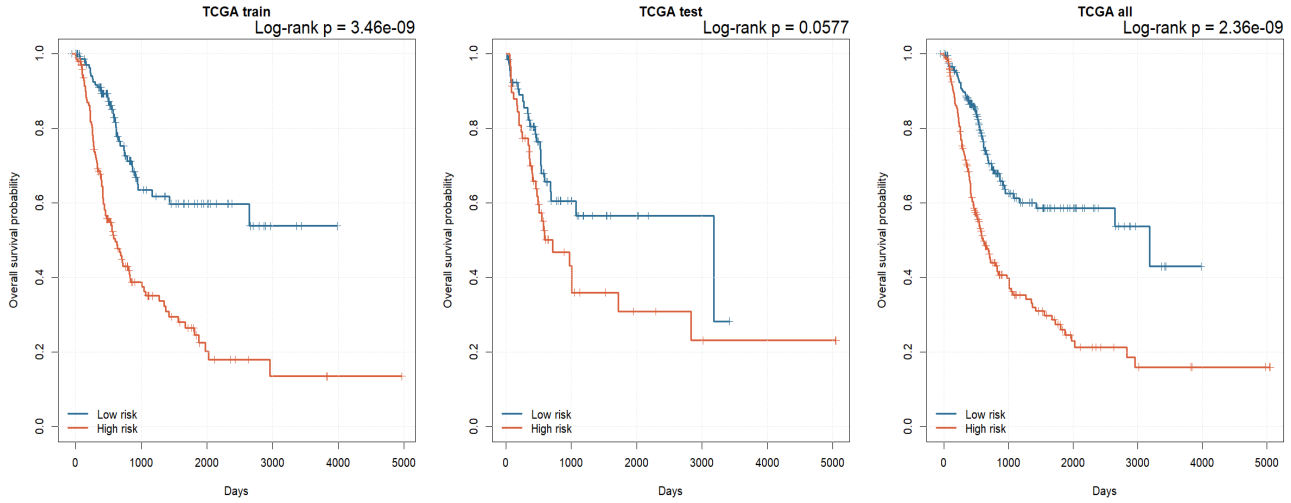


*Supplementary Figure S15. Distribution of the fixed five-gene risk score across all TCGA-BLCA tumor samples.*

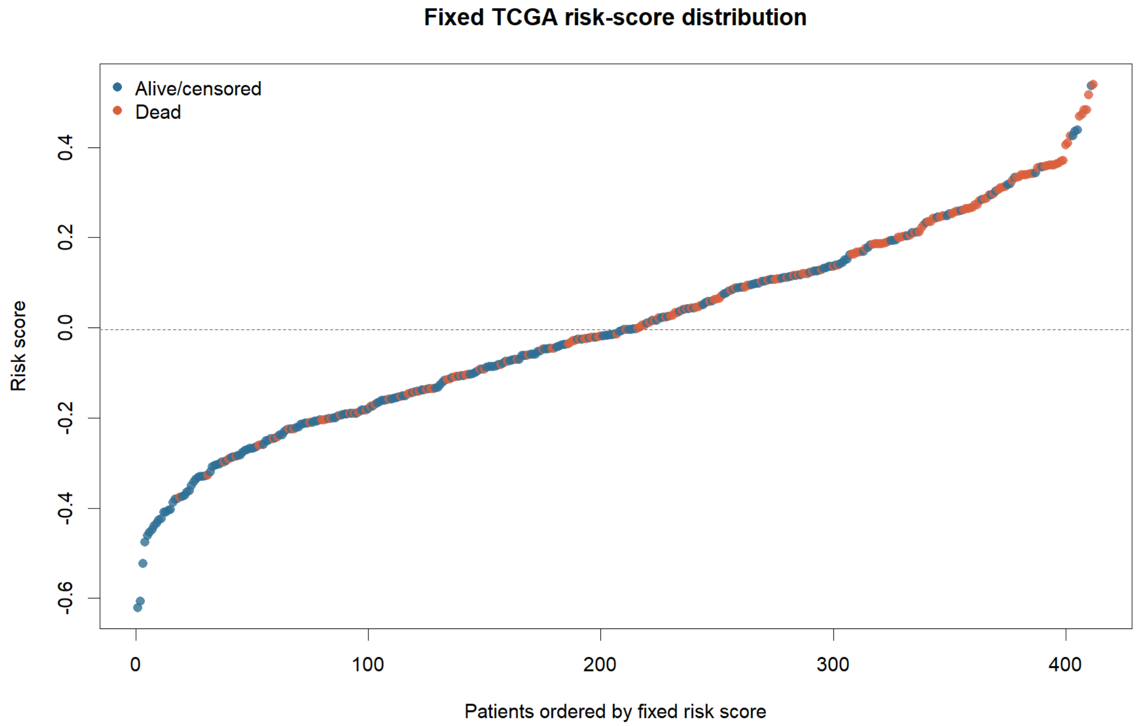

*Supplementary Figure S16. Time-dependent ROC curves for the fixed five-gene model in the full TCGA-BLCA cohort.*

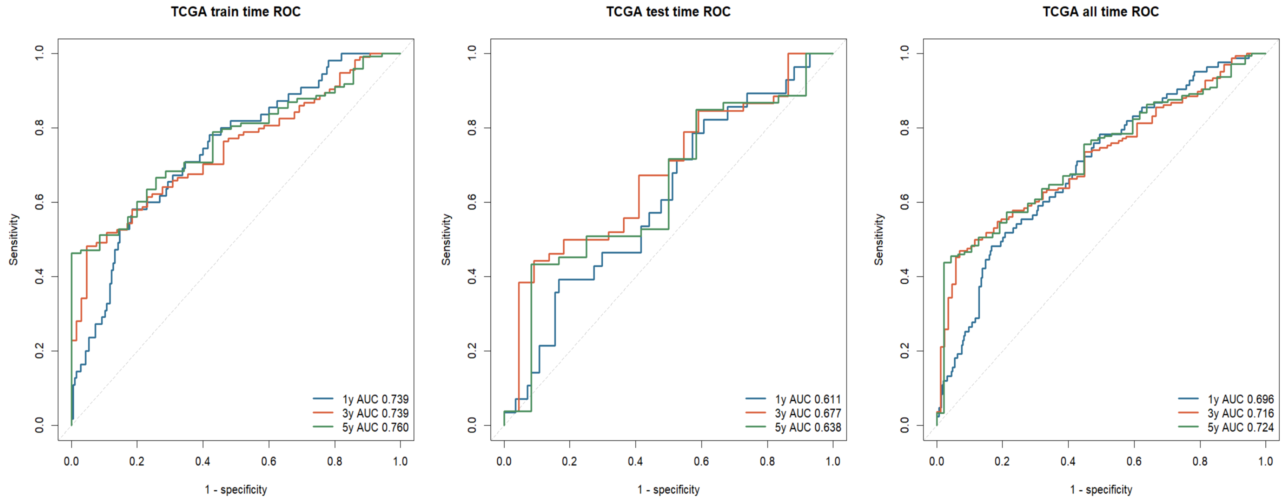


*Supplementary Figure S17. Kaplan-Meier overall survival curves for the fixed five-gene score in the external GEO validation cohorts GSE13507 and GSE31684, stratified by median risk score.*

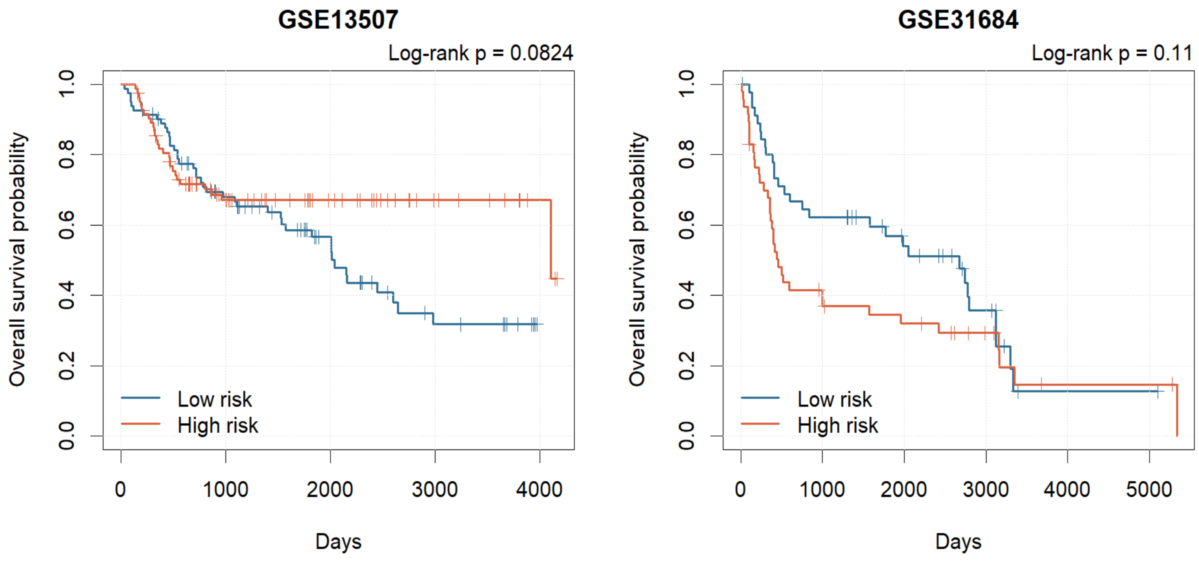

*Supplementary Figure S18. Distribution of the fixed five-gene risk score in the GSE13507 and GSE31684 validation cohorts.*

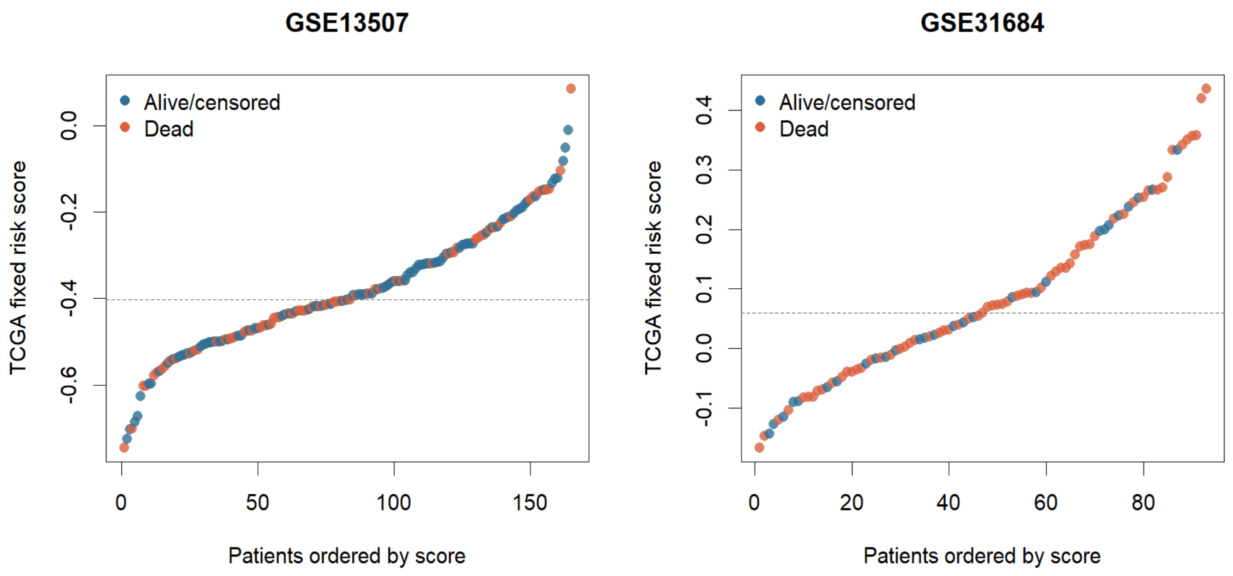


*Supplementary Figure S19. Time-dependent receiver operating characteristic (ROC) curves for the fixed five-gene score in the GEO validation cohorts.*

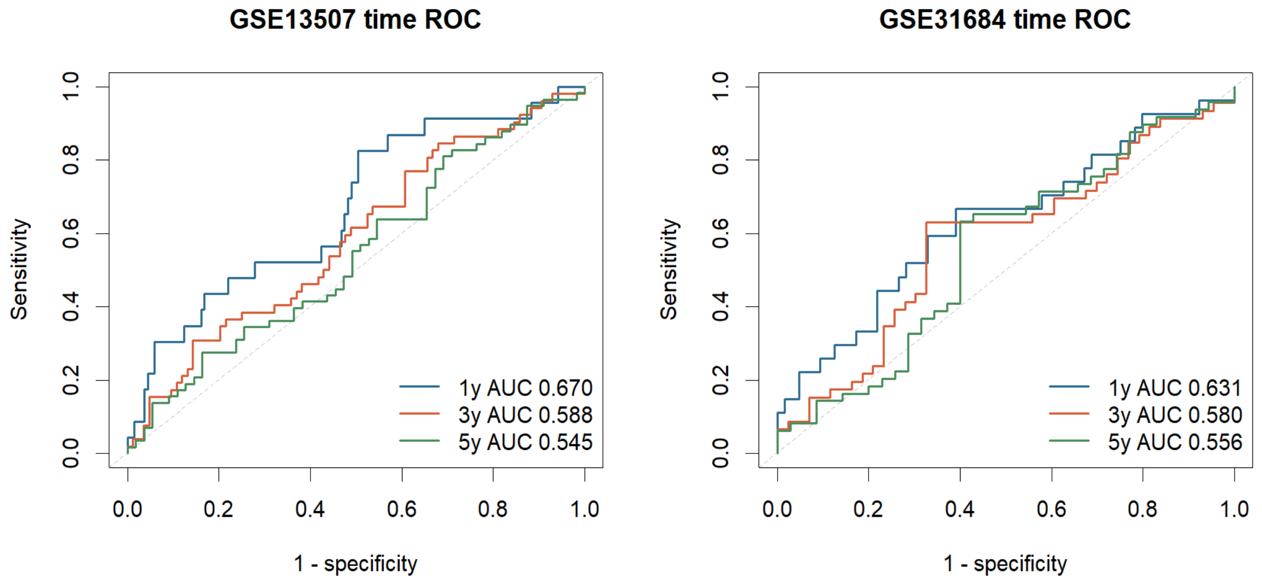

*Supplementary Figure S20. Distribution of the fixed five-gene risk score in the GSE13507 and GSE31684 cohorts after applying the reduced model formula without refitting.*

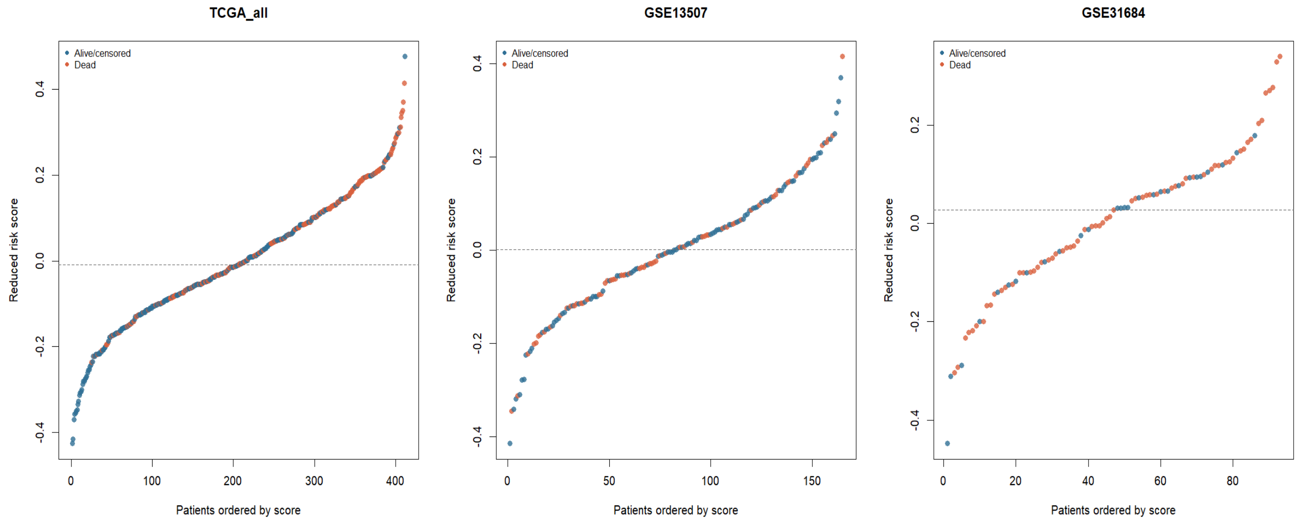

*Supplementary Figure S21. Tumor-normal differential expression landscape in TCGA-BLCA, showing the exploratory Wilcoxon screen used to support biological interpretation and figure generation.*

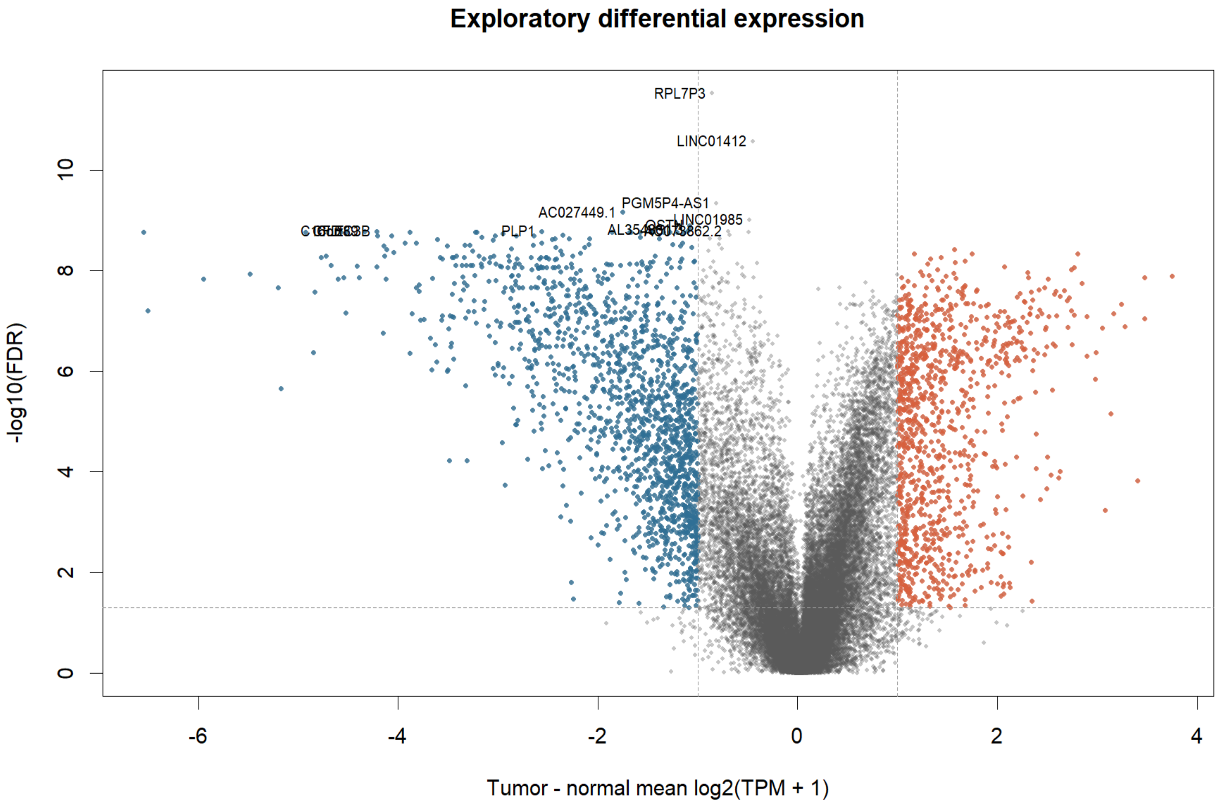

*Supplementary Figure S22. Heatmap of the top 50 differentially expressed genes between tumor and adjacent normal bladder tissue, hierarchically clustered by sample.*

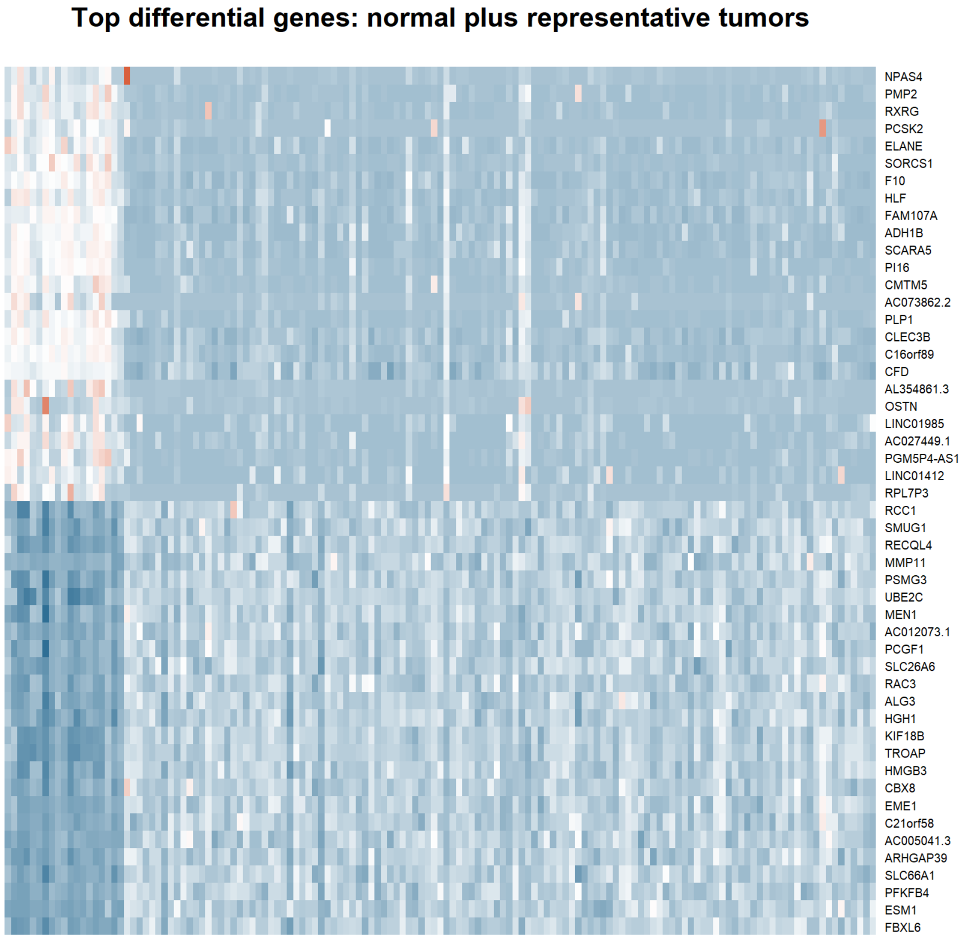

*Supplementary Figure S23. Bar plot of the top differentially expressed genes ranked by absolute log2 fold change between tumor and normal samples.*

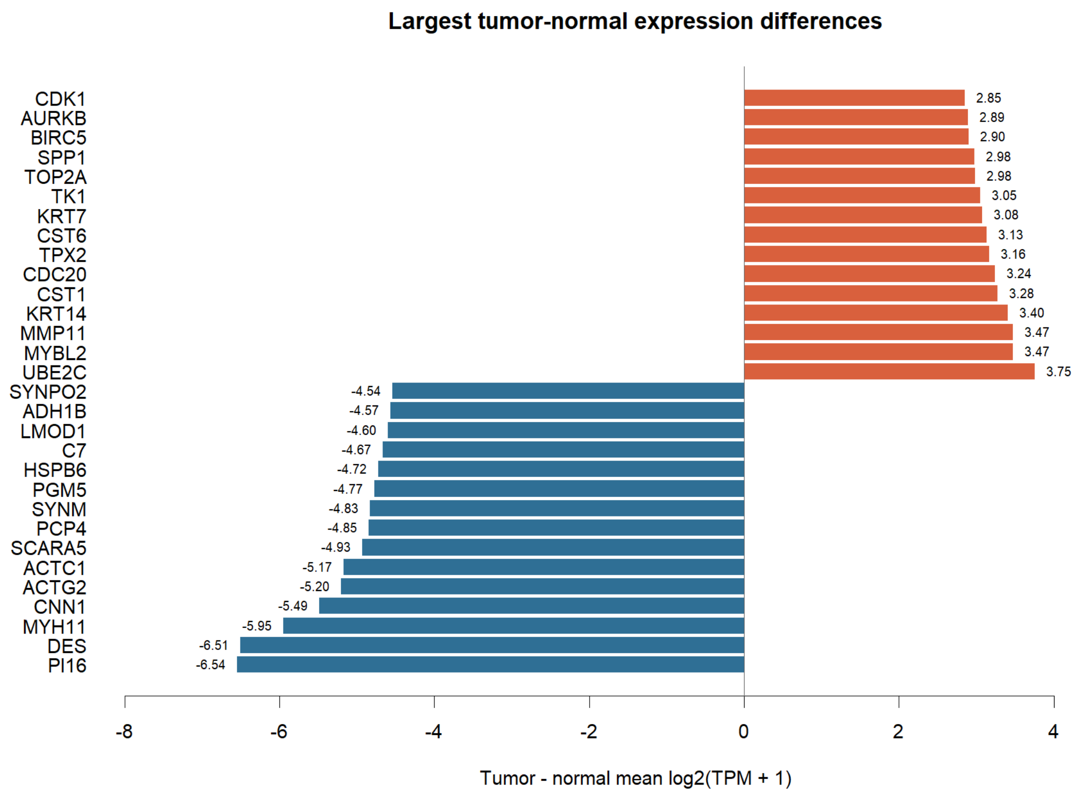

*Supplementary Figure S24. Bar plot of the top exploratory tumor-versus-normal genes ranked by log2 fold change, supporting the differential expression screen described in the Methods.*

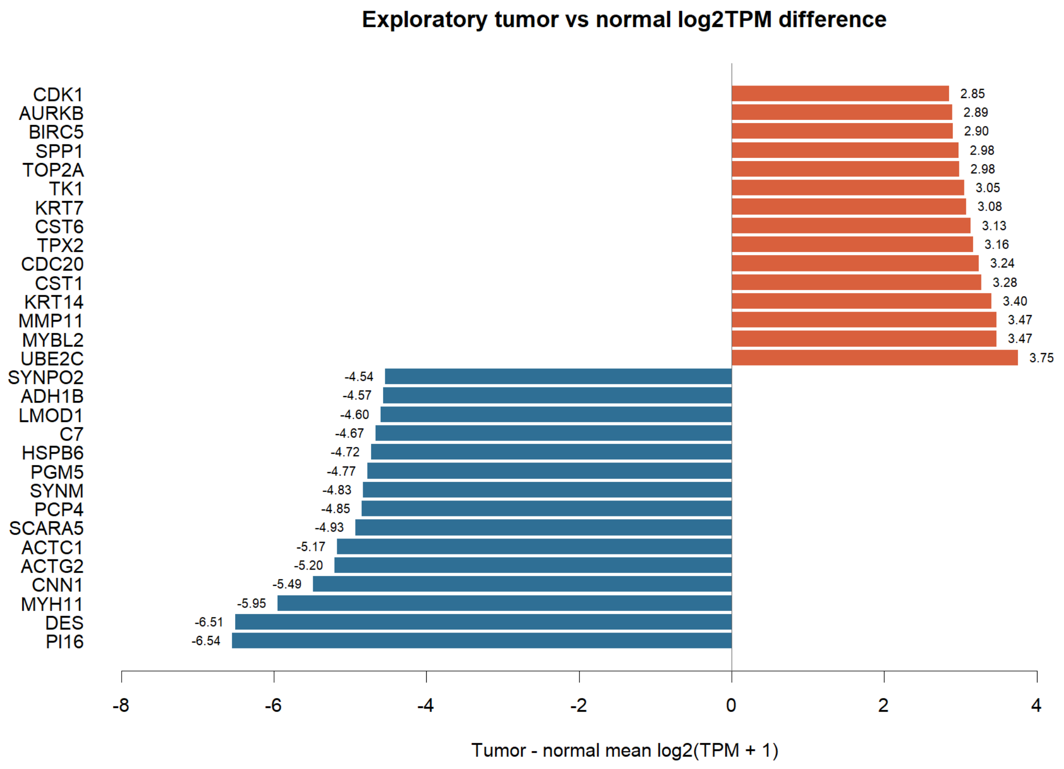

*Supplementary Figure S25. Volcano plot of differentially expressed genes between high-risk and low-risk TCGA tumors defined by the fixed five-gene score.*

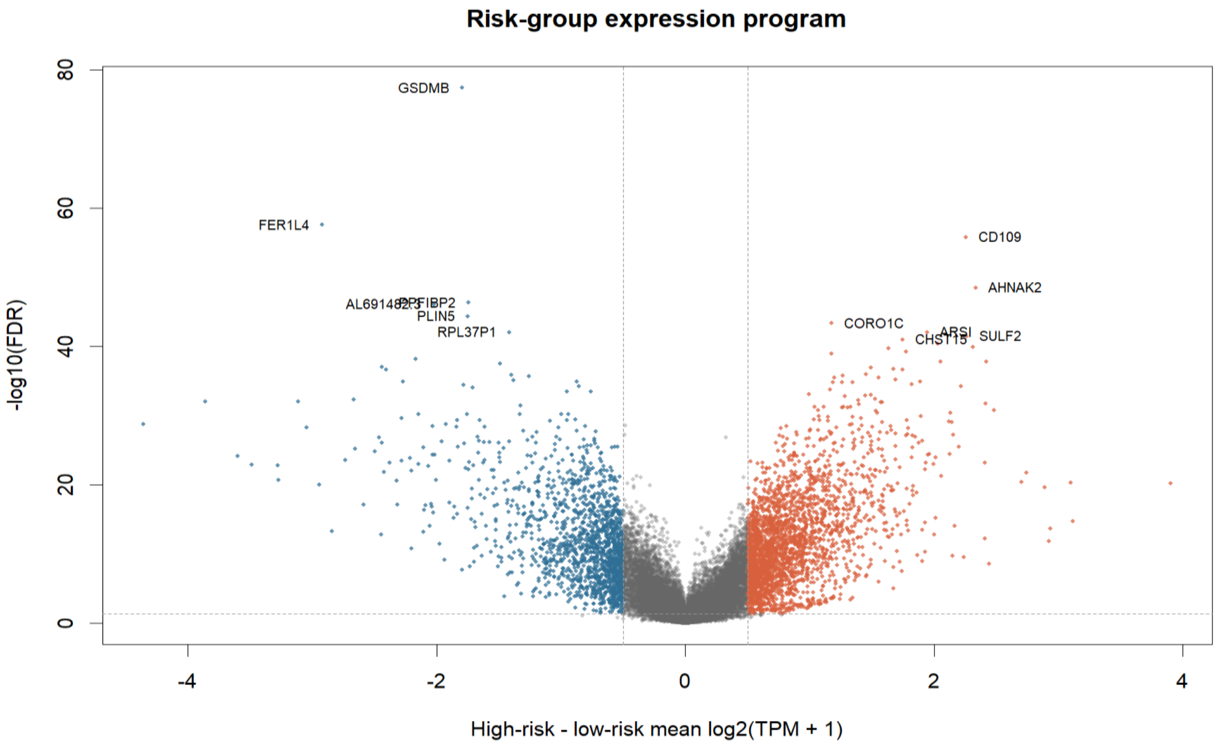

*Supplementary Figure S26. Boxplots of immune and stromal signature scores (including CAF, macrophage, checkpoint, cytotoxic/CD8, and inflammatory programs) compared between high-risk and low-risk TCGA tumors.*

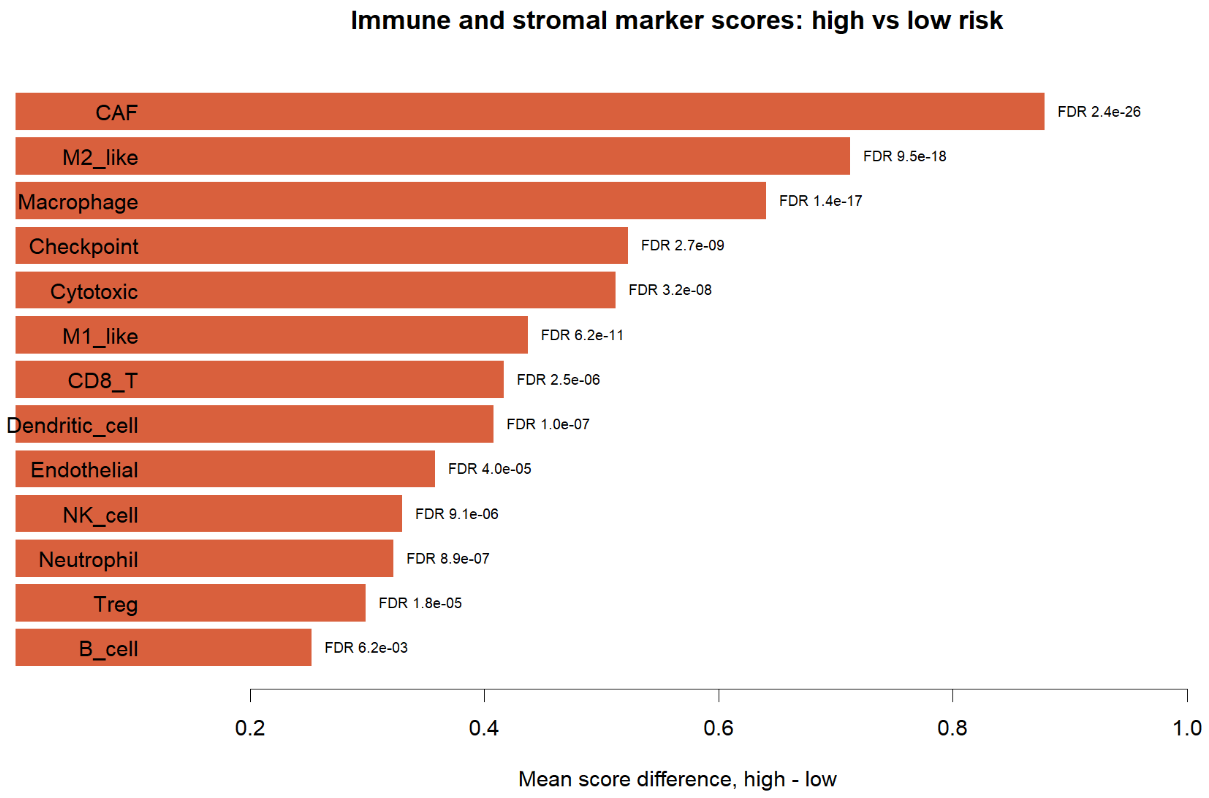

*Supplementary Figure S27. Distribution of nonsilent somatic mutation burden in TCGA-BLCA tumors, compared between high-risk and low-risk groups.*

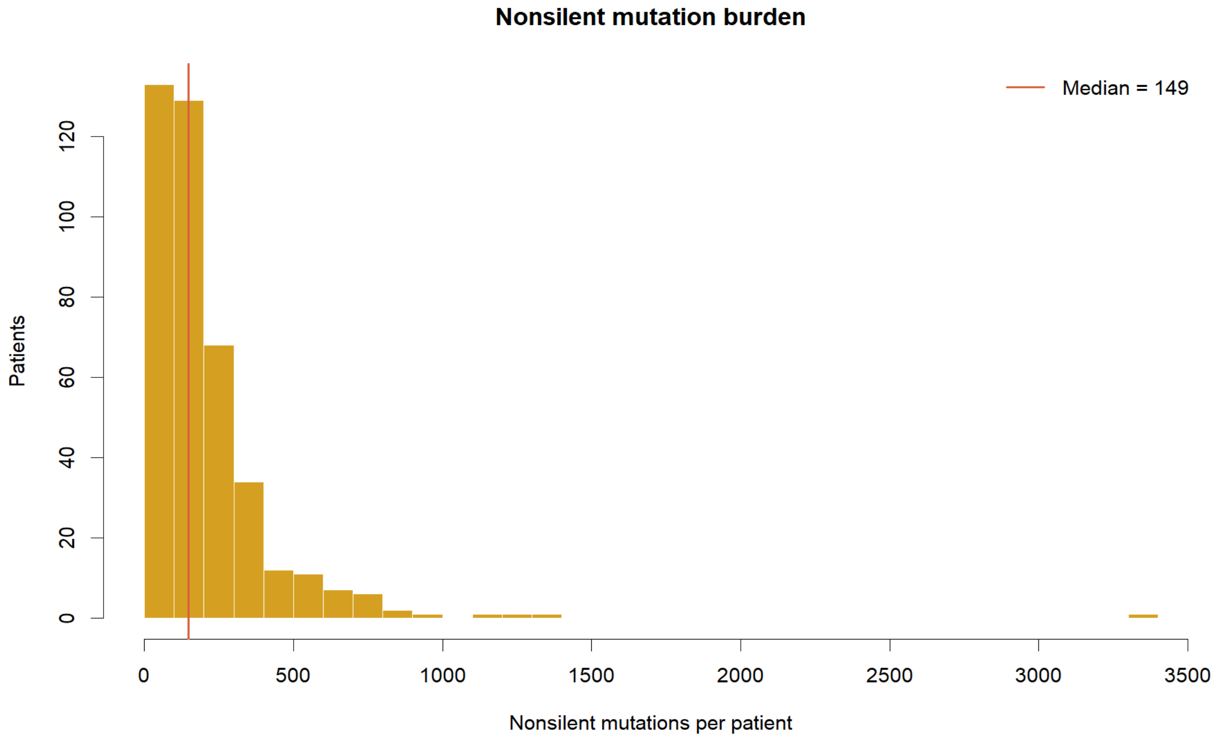

*Supplementary Figure S28. Heatmap of the most variable RPPA protein features, compared between high-risk and low-risk TCGA tumors.*

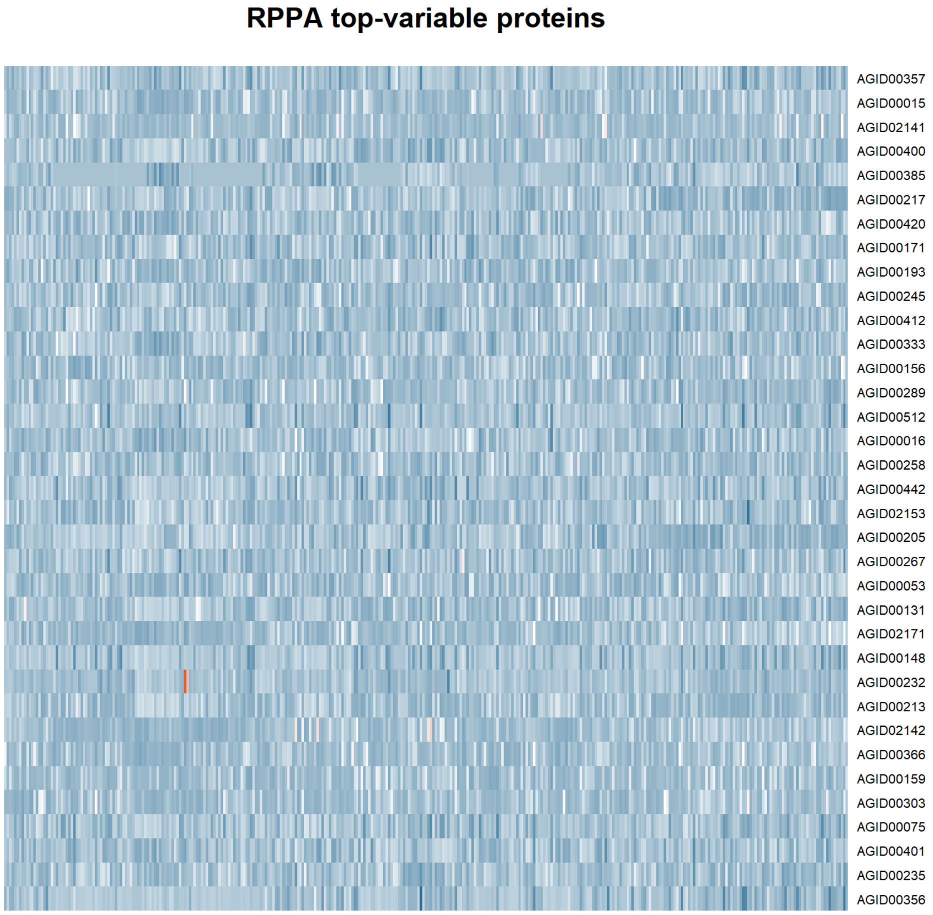

*Supplementary Figure S29. Copy-number variation burden (mean absolute segment amplitude and total altered length) compared between high-risk and low-risk TCGA tumors.*

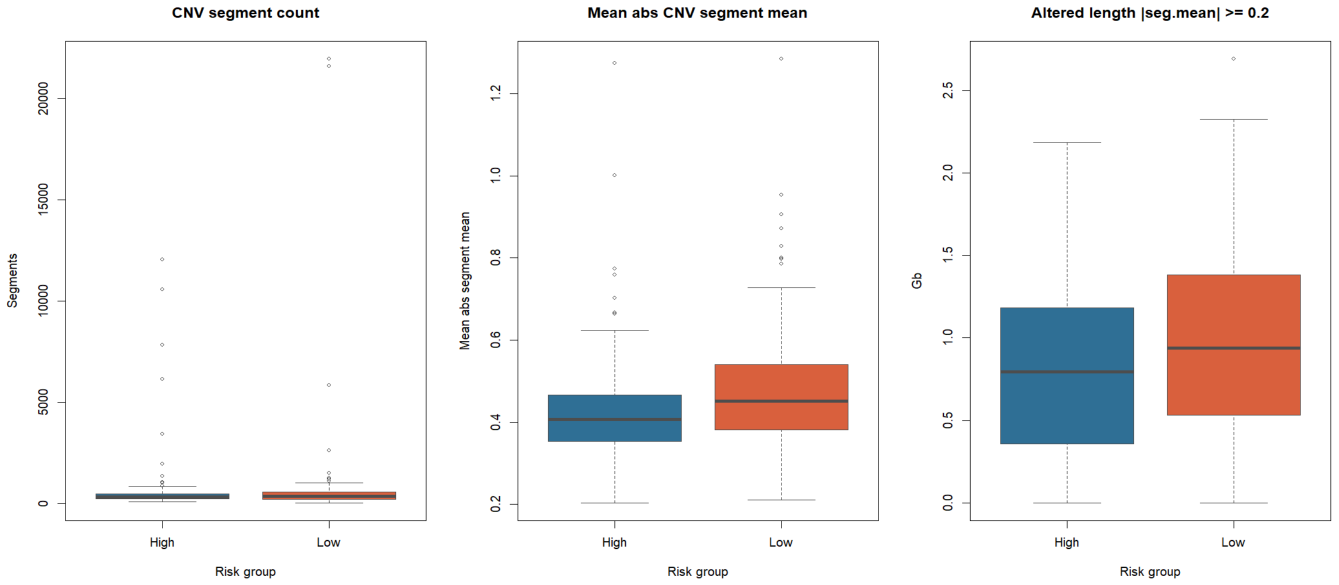


*Supplementary Figure S30. Comparison of RPPA protein expression features between high-risk and low-risk TCGA tumors.*

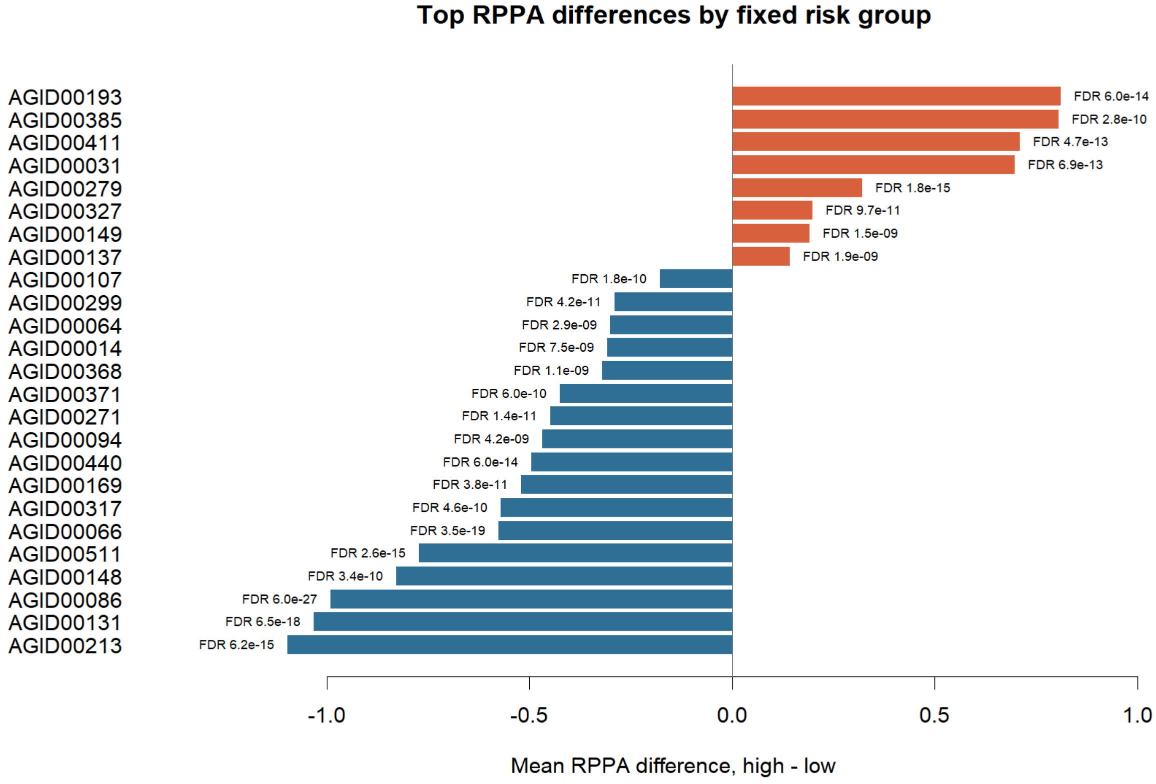

*Supplementary Figure S31. Bar plot of the most frequently mutated genes in the TCGA-BLCA cohort.*

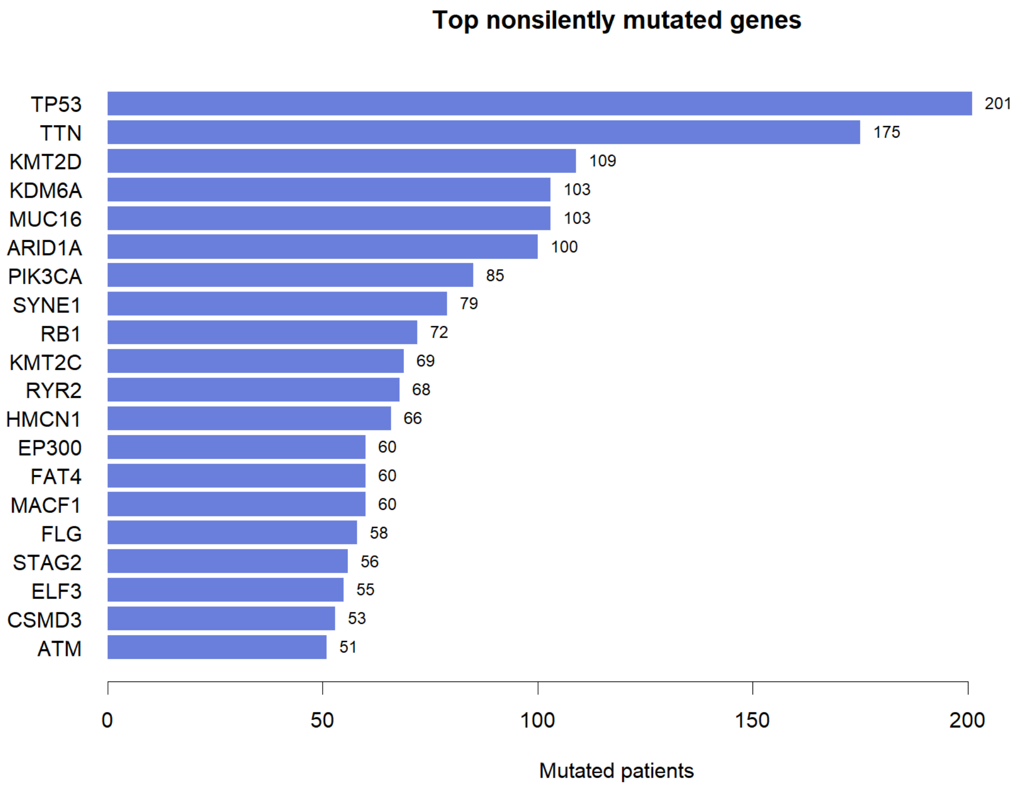

*Supplementary Figure S32. Cell-type-resolved risk-score distribution in the candidate single-cell validation datasets (GSE222315 and GSE293189). Dot size and color represent the proportion and mean fixed-model risk score of cells in each annotated compartment, showing the highest scores in endothelial and fibroblast populations.*

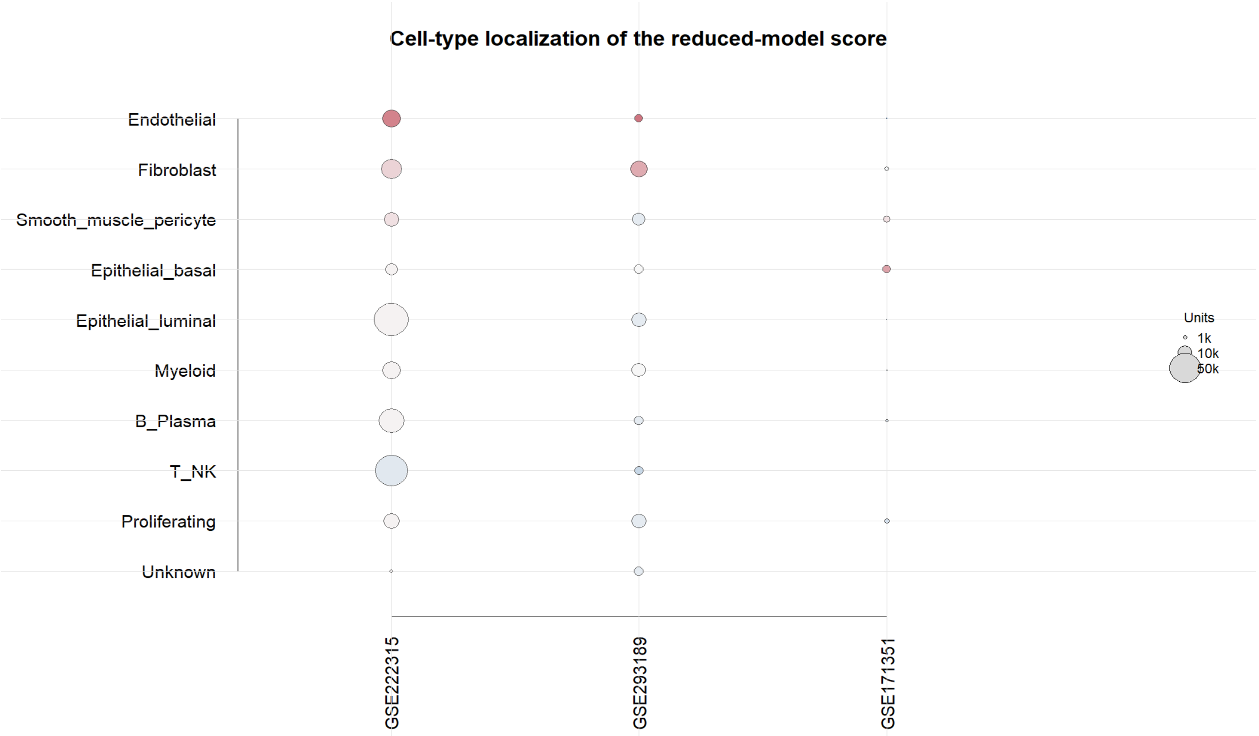

*Supplementary Figure S33. Heatmap summary of mean risk score by cell type across the candidate single-cell validation datasets, corresponding to the dotplot in Supplementary Figure S2 and confirming the endothelial/fibroblast/stromal-dominant pattern observed in GSE145137.*

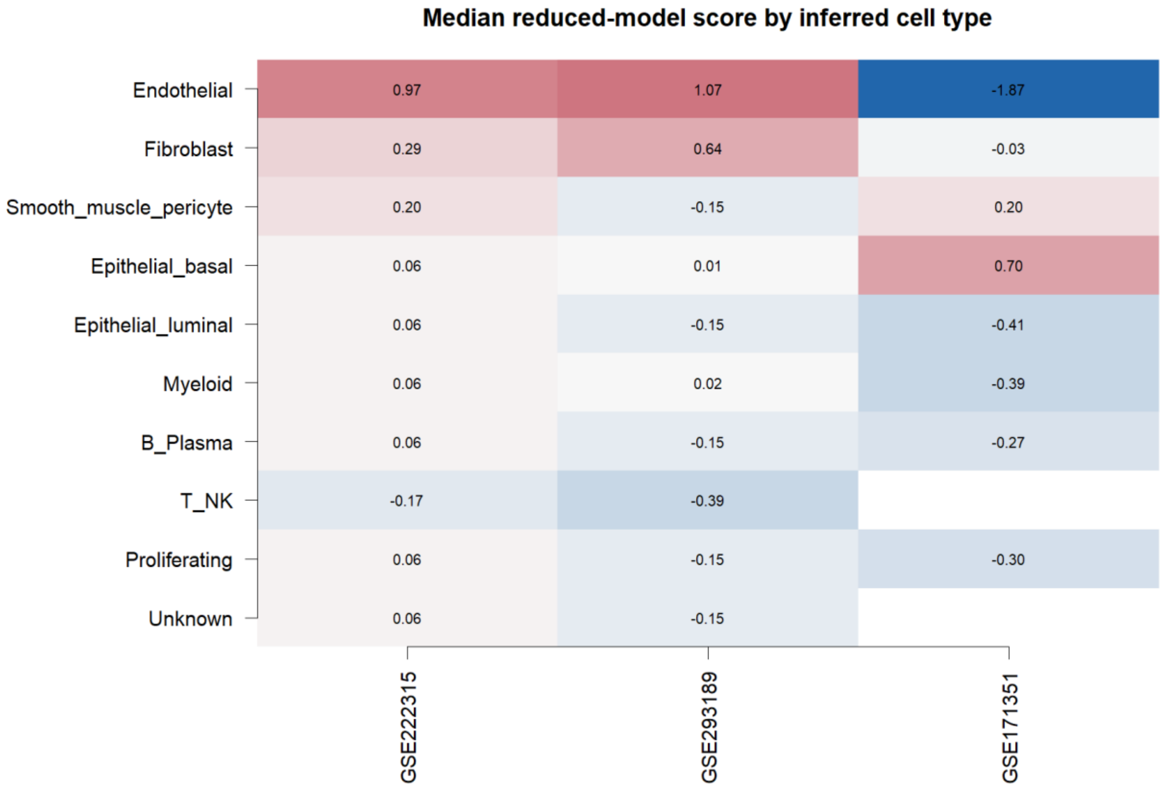

*Supplementary Figure S34. Cell-type composition of the GSE145137 primary bladder cancer single-cell RNA-seq sample, based on author-provided annotations.*

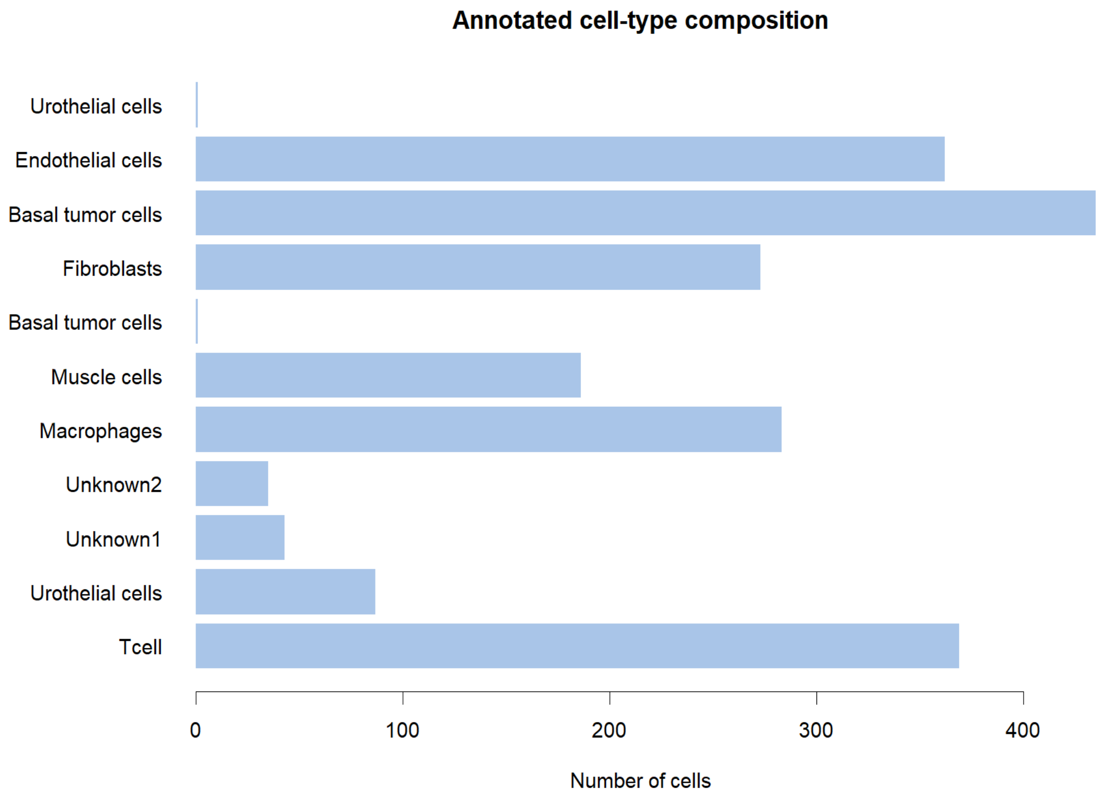

*Supplementary Figure S35. Heatmap of the contribution of each of the five fixed-model genes to the risk score across annotated cell types in GSE145137.*

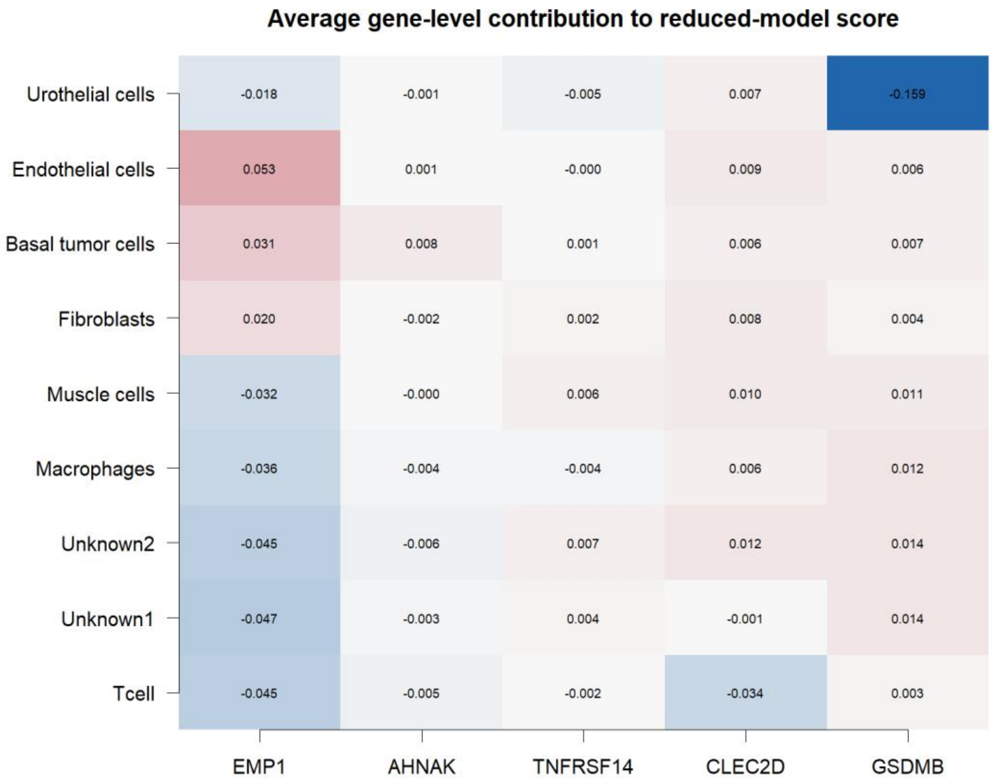

*Supplementary Figure S36. Dotplot of the five fixed-model gene expression levels across annotated cell types in the GSE145137 single-cell dataset.*

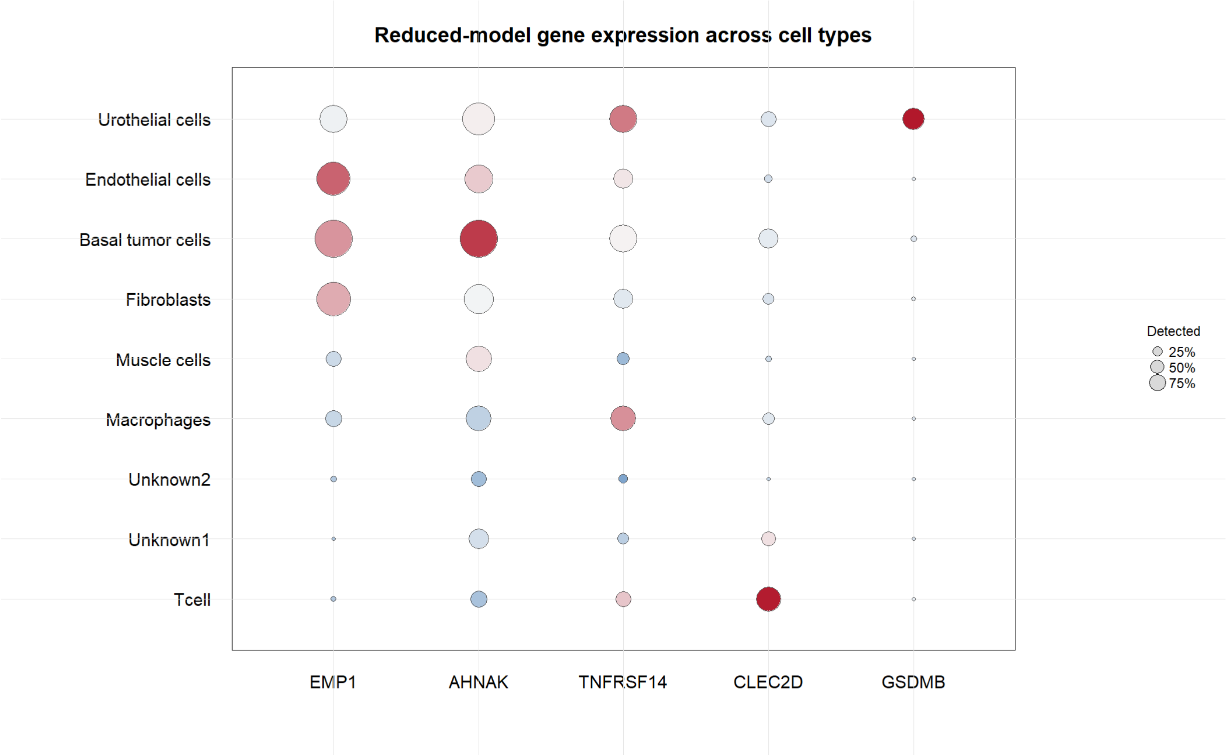

*Supplementary Figure S37. PCA/UMAP projection of GSE145137 single cells, colored by author-provided cell-type annotation.*

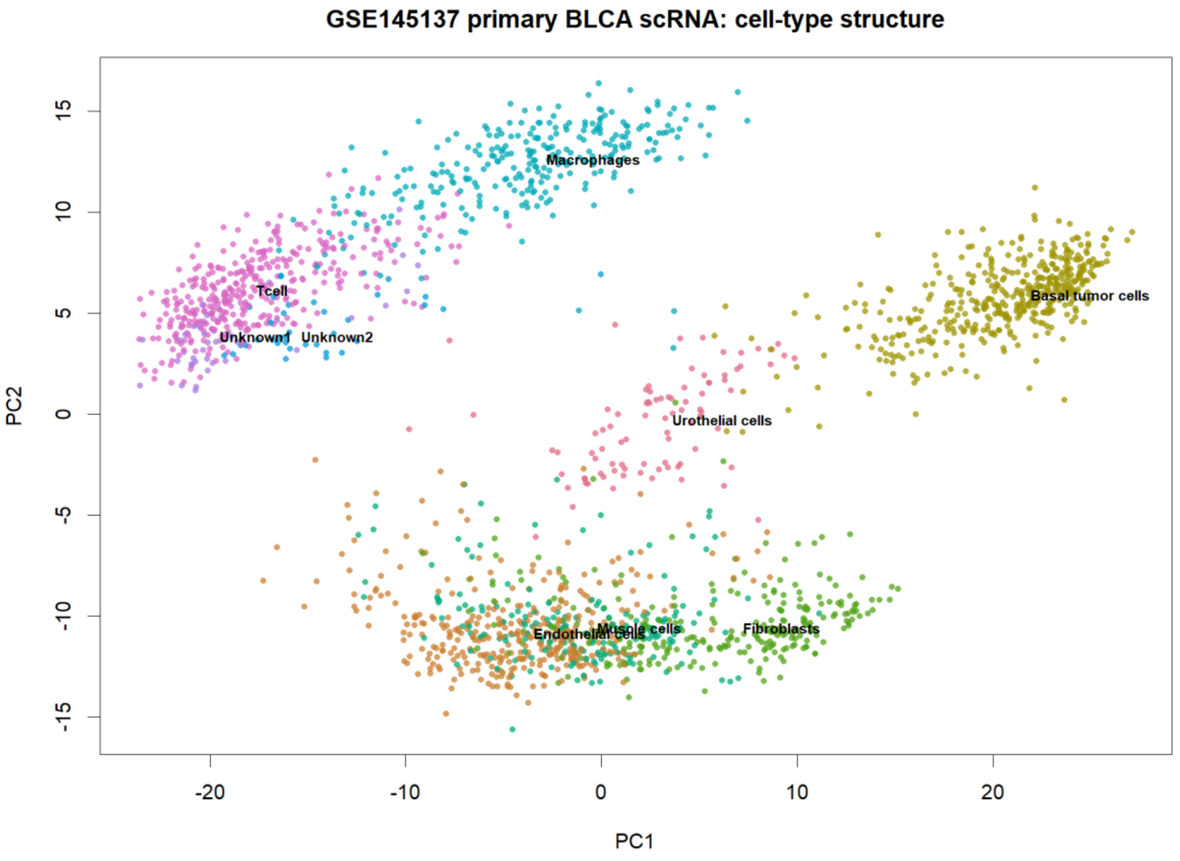

*Supplementary Figure S38. PCA/UMAP projection of GSE145137 single cells, colored by fixed five-gene risk score, showing enrichment in basal tumor, endothelial, and fibroblast compartments.*

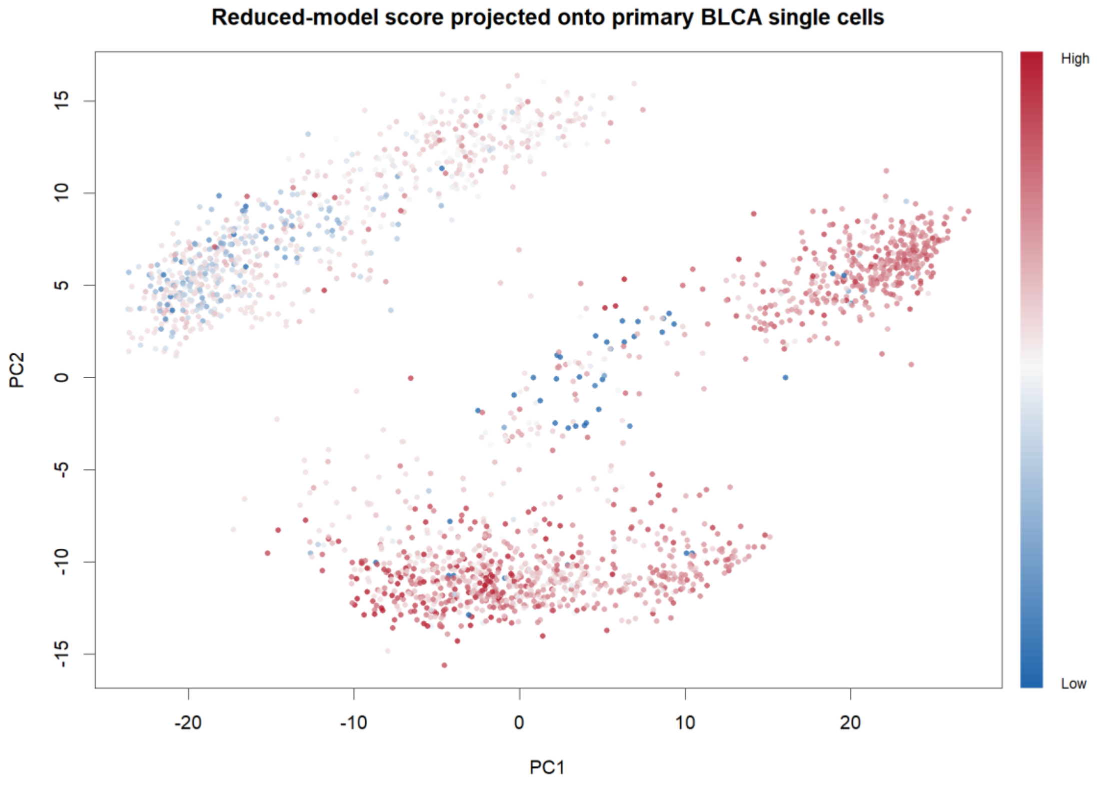

*Supplementary Figure S39. Immunotherapy response validation using the fixed risk score in ICB-treated cohorts, showing time-dependent ROC curves for risk score predicting non-response in IMvigor210 and GSE176307.*

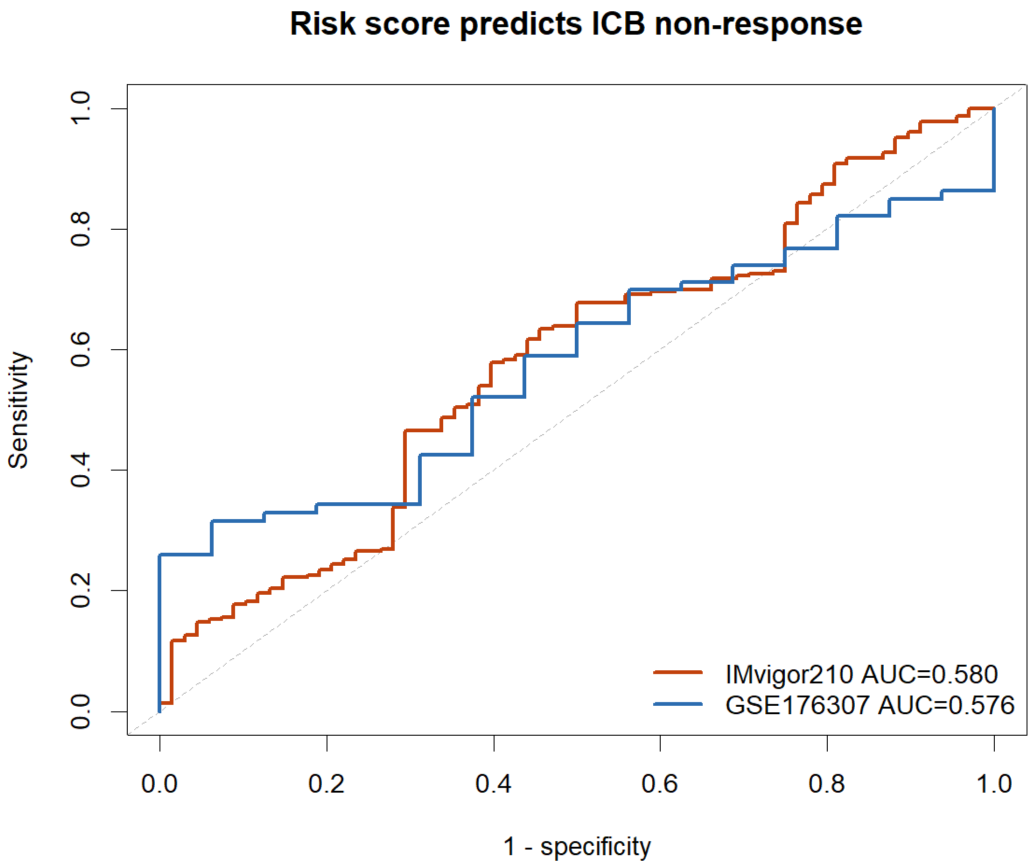

*Supplementary Figure S40. Balance between cytotoxic/CD8 effector signatures and TGF-beta/CAF suppression signatures by risk group in the ICB-treated cohorts, illustrating the immune-present but suppressed phenotype of high-risk tumors.*

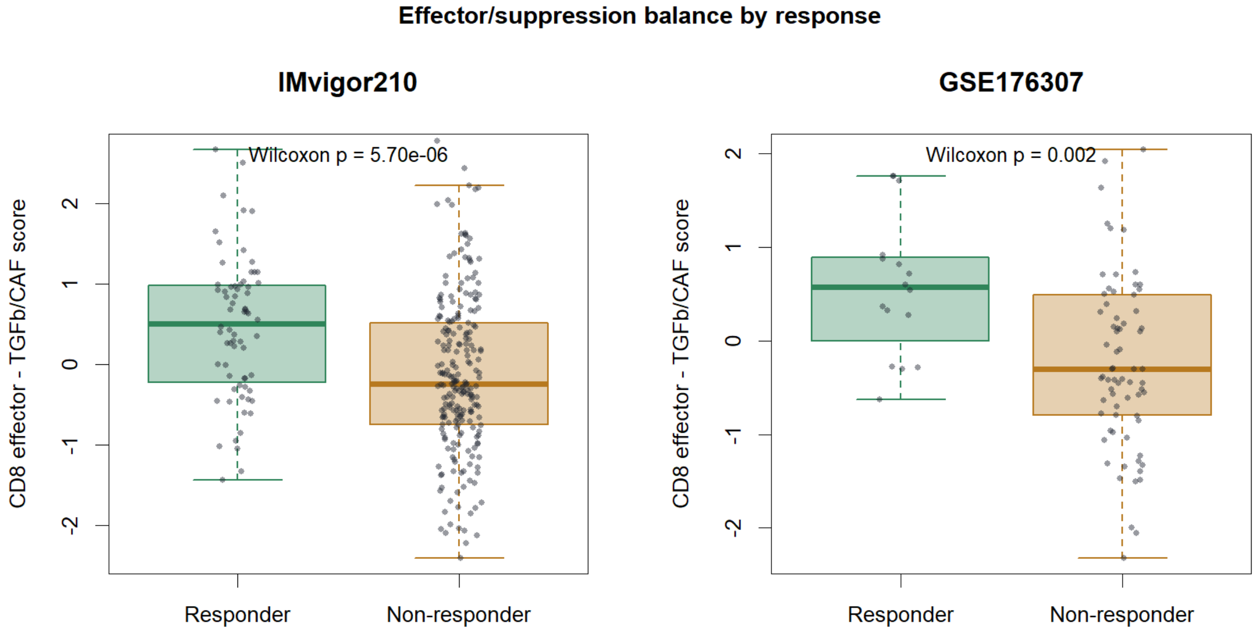


*Supplementary Figure S41. Kaplan-Meier overall survival curves by median-split risk group in the IMvigor210 and GSE176307 immune-checkpoint-blockade cohorts.*

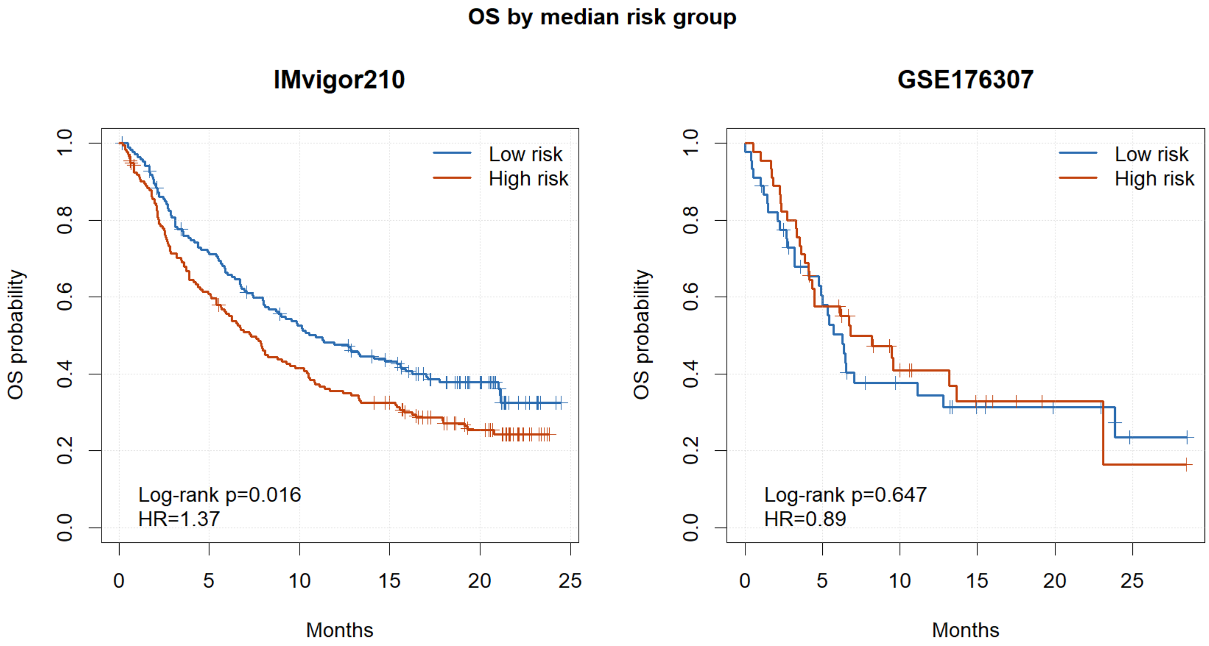

*Supplementary Figure S42. Kaplan-Meier progression-free survival curves by median-split risk group in the ICB-treated cohorts.*

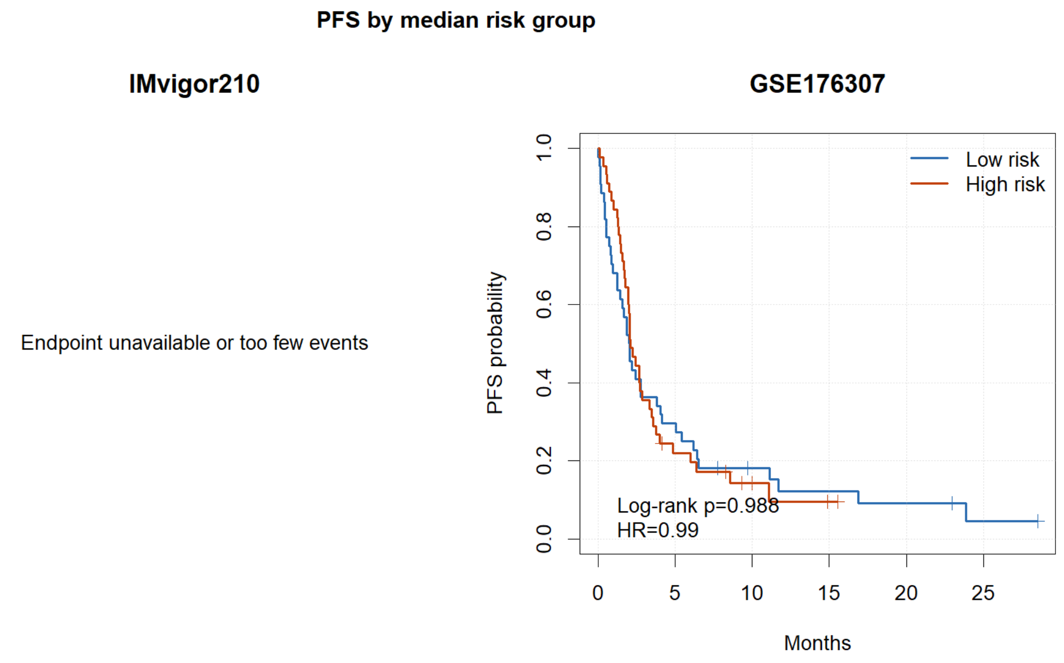


*Supplementary Figure S43. Heatmap of the five fixed-model gene expression values across samples in the IMvigor210 and GSE176307 cohorts, ordered by risk group and response status.*

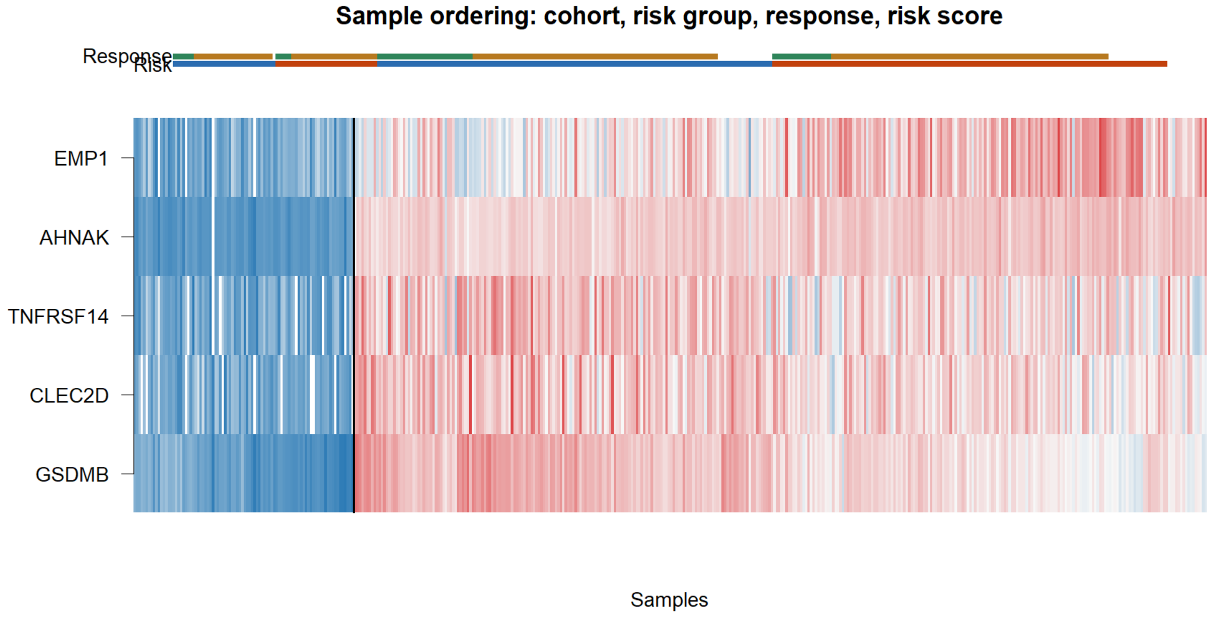

*Supplementary Figure S44. Boxplot comparison of the fixed five-gene risk score between immunotherapy responders (CR/PR) and non-responders (SD/PD) in the ICB-treated cohorts.*

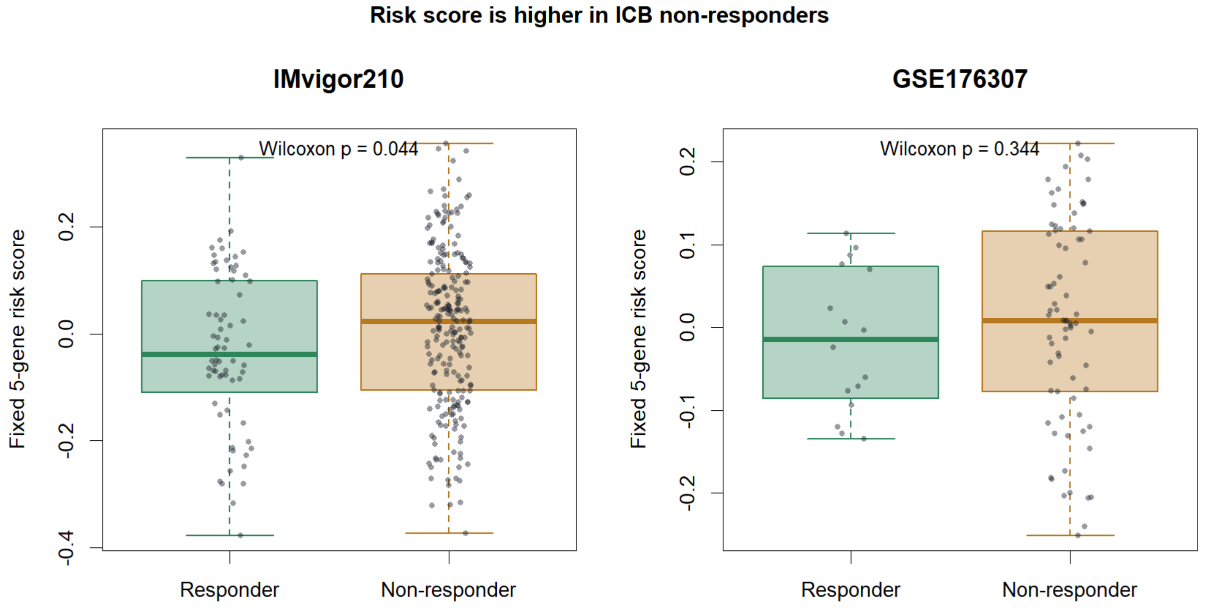


*Supplementary Figure S45. Stacked bar plot of immunotherapy response rate (responder versus non-responder) by median-split risk group in IMvigor210 and GSE176307.*

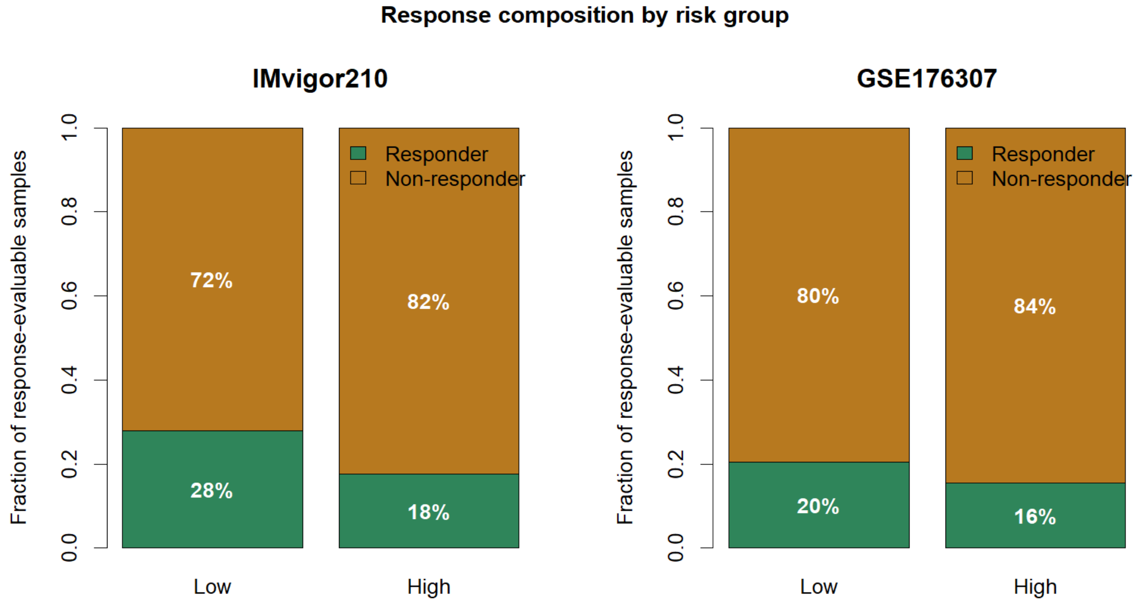

*Supplementary Figure S46. Correlation heatmap between the fixed five-gene risk score and immune/stromal signature scores (including TGF-beta/CAF, EMT-stromal, luminal epithelial, and CD8/checkpoint programs) in the IMvigor210 and GSE176307 cohorts.*

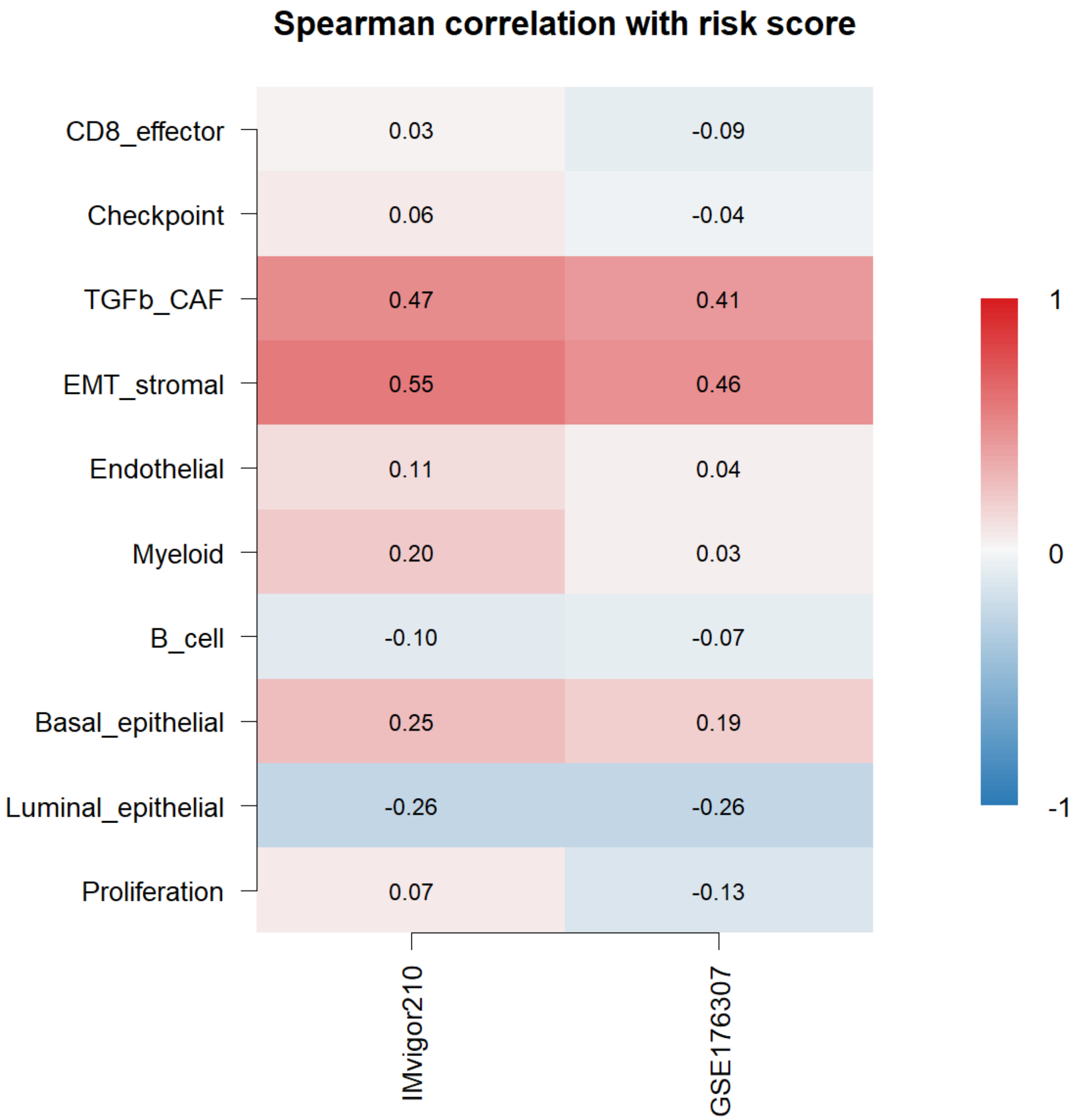
